# Environmental exposure to antibiotics drives and sustains antibiotic resistance in *Neisseria gonorrhoeae*: A modelling study

**DOI:** 10.64898/2025.12.08.692580

**Authors:** Ernst D. Schäfer, Constance Schultsz, Henry J. C. de Vries, Alje P. van Dam, Rik Oldenkamp

## Abstract

Ciprofloxacin resistance is widespread in *Neisseria gonorrhoeae*, despite its discontinuation as recommended treatment option in 2007. Unintentional low-level exposure to ciprofloxacin, e.g., environmental or food-borne, might provide sufficiently high selection pressure to maintain resistance and compensate potential fitness costs. We used a mathematical within-host model of gonorrhoea to investigate the effect of low-level ciprofloxacin exposure on susceptibility. We present a mechanistic within-host model of gonococcal infections in the male urethra, to simulate the influence of low-level ciprofloxacin exposures on gonococcal ciprofloxacin susceptibility. We assessed the impact of repeated exposures to varying amounts of ciprofloxacin 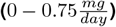 in a chain of sequentially transmitted infections. Daily exposure to at least 0.15*mg* ciprofloxacin was sufficient to select for low-level resistance. Moreover, in the absence of ciprofloxacin treatment, resistant variants could revert back to variants with lower levels of resistance. However, background exposure can balance fitness costs of resistance and prevent a complete reversion to the fully susceptible variant. Even for human-host restricted pathogens such as *Neisseria gonorrhoeae*, a One Health approach that considers unintentional antibiotic exposures is needed to avoid the emergence and spread of resistance.

## Introduction

Gonorhoea is one of the most prevalent sexually transmitted bacterial infections, with an estimated 78.3 million new infections every year, and its incidence is on the rise in many countries Bodie et al. (2019). Gonococcal resistance to previous first-line antibiotics is widespread and the emergence of strains resistant to current antibiotics has necessitated the development of new treatment options, improvement of surveillance, diagnostics and prevention, and the development and implementa-tion of national and global response plans Bodie et al. (2019).

Since the introduction of antibiotics for gonorrhoea treatment, resistant strains have emerged and spread Costa-Lourenço et al. (2017). WHO guidelines recommend changing first-line antibiotic treatment 5% prevalence. In the subsequent absence of selection pressure, fitness costs associated with resistance to the abandoned antibiotic should lead to resistant strains being outcompeted by susceptible strains or strains resistant to the replacement antibiotic. In the 1980s, fluoroquinolones, mainly ciprofloxacin, were introduced as treatment of gonorrhoea, and ciprofloxacin was abandoned as first-line treatment in the 2000s. However, since then gonococcal resistance to ciprofloxacin increased in Australia, China, and the USA Hooshiar et al. (2024). A recent phylodynamic analysis of N. gonorrhoea in the USA Helekal et al. (2025) indeed indicated that whilst in most fluoroquinolone-resistant lineages resistance-conferring alleles were replaced with wild-type alleles after the abandonment of ciprofloxacin, resulting in phenotypic susceptibility, gonococcal strains carrying GyrA 91F/95A mutations appeared to have a fitness advantage over susceptible wildtype strains, even when ciprofloxacin was no longer in use Helekal et al. (2025). The authors proposed bystander exposure as one possible explanation, where N. gonorrhoeae is exposed to ciprofloxacin that is intended to treat another (co)infection Olesen and Grad (2020). This hypothesis is at least partly supported by a large scale ecological analysis through which positive correlations were found between the prevalence of gonococcal resistance to ciprofloxacin and country-level fluoroquinolone consumption Kenyon et al. (2020).

Ciprofloxacin concentrations much lower than therapeutic concentrations 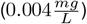 can already select for resistance in *N. gonorrhoeae in vitro* González et al.(2022). As such, resistance might be selected for, and maintained by, unintentional exposures to low concentrations of ciprofloxacin. The presence of antibiotics, including quinolones, has been demonstrated in surface waters, drinking waters, animal and vegetable food products, and soil and air particles Ben et al. (2019). These environmental reservoirs of antibiotics have long been recognised as potential drivers of antibiotic resistance through the selection pressure they induce on human and natural microbial systems Ben et al. (2019). To test our hypothesis that repeated low-level ciprofloxacin exposures can select for ciprofloxacin resistance in *N. gonorrhoeae*, we used a mechanistic modelling approach. Within-host mathematical models aim to capture the dynamics of an infection through differential equations governing cellular, bacterial, and/or viral populations. By integrating a within-host model of *N. gonorrhoeae* infection Jayasundara et al. (2019) with pharmacokinetic and pharmacodynamic models, the emergence of and selection for resistance determinants can be simulated Witzany et al. (2023).

## Methods

WIGWAM (Within-host Infection dynamics of neisseria Gonnorhoeae With Antibiotic resistance Modelling) simulates the dynamics of male urethral *N. gonorrhoeae* infection. The model iterates on the model developed and presented by Jayasundara et al. Jayasundara et al. (2019), with major differences in the representation of host-pathogen interactions. The WIGWAM source code is available at https://gitlab.com/aighd-modelling/WIGWAM.

After *N. gonorrhoeae* is transmitted, it colonises and invades epithelial mucosal tissues, using opacity-associated (Opa) proteins and type IV pili Quillin and Seifert (2018). After endocytosis, gonococci can replicate within epithelial cells. *N. gonorrhoeae* is able to partially evade both oxidative and non-oxidative killing mechanisms employed by the neutrophils recruited to the site of infection upon gonococcal colonisation Palmer and Criss (2018). Moreover, phagocytosed gonococci can survive and reproduce inside neutrophils and macrophages Quillin and Seifert (2018). In urethral infections, a common symptom is the discharge of a neutrophil-rich exudate, which facilitates bacterial transmission to new hosts Quillin and Seifert (2018).

The absence of an adaptive immune response to gonorrhoea, and thus the lack of long term immunity, makes reinfections common. However, despite the suppression of the adaptive immune response, IgM producing memory B cells are activated by gonorrhoea in a T-cell-independent response So et al. (2012).

WIGWAM simulates the following interacting populations: unattached extracellular bacteria (*B*_*u*_), extracellular bacteria attached to the epithelium (*B*_*a*_), bacteria inside the epithelium (*B*_*i*_), bacteria surviving within neutrophils (*B*_*s*_), neutrophils (*N*), immune signalling cells such as T helper lymphocytes (*L*), and IgM-producing memory B cells (*M*). Figure 1 provides a schematic overview of the model interactions.

**Figure 1.**
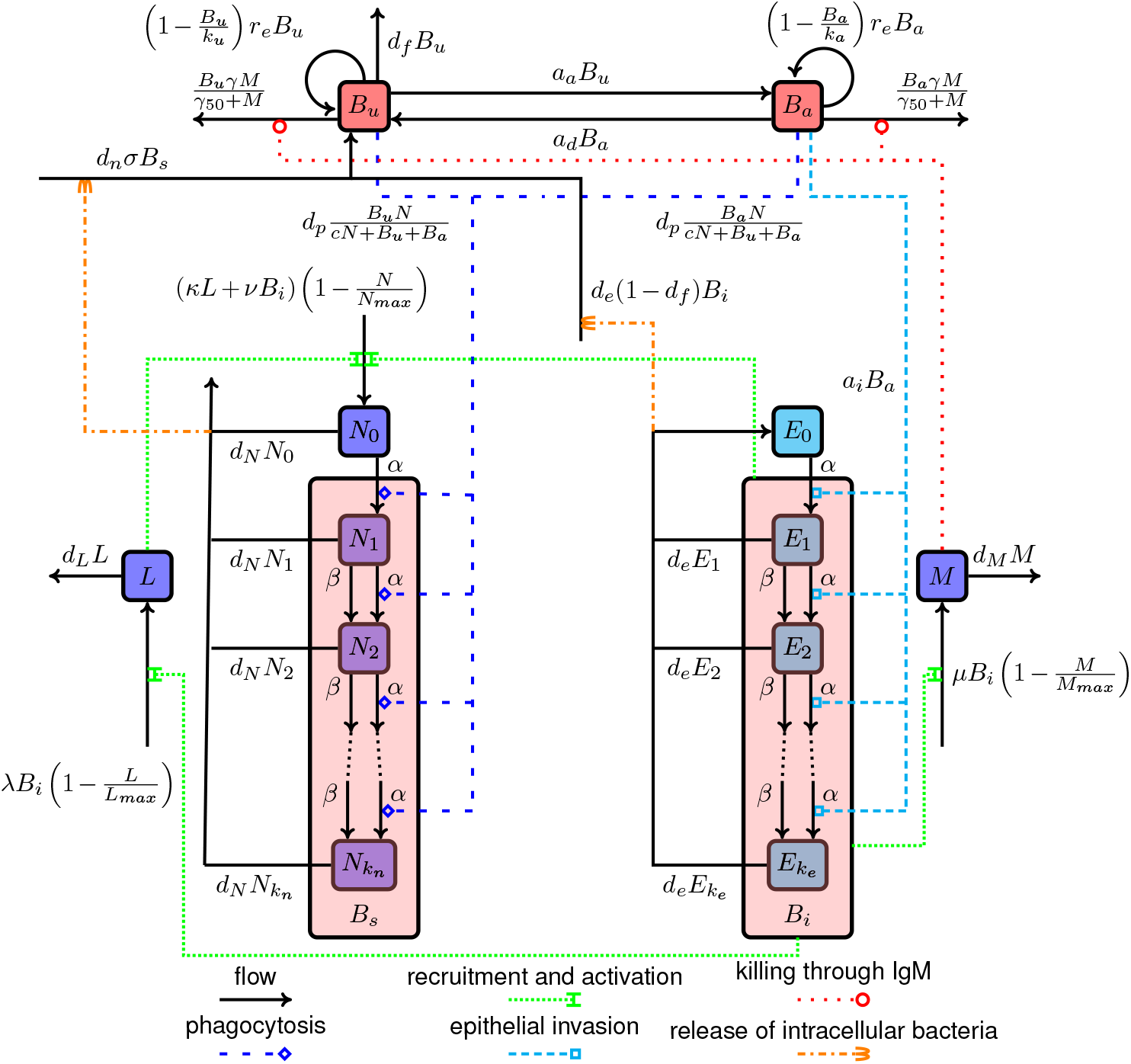
Schematic representation of the WIGWAM model. Here *B*_*u*_ = unattached bacteria, *B*_*a*_ = attached bacteria, *B*_*i*_ = bacteria inside the epithelium, *B*_*s*_ = bacteria surviving in neutrophils, *L* = signalling lymphocytes, *M* = memory B cells, *N*_*x*_ = neutrophils which contain *x* bacteria, *E*_*x*_ = epithelial cells which contain *x* bacteria, *α* = group change due to bacteria entering host cell, *β* = group change due to bacteria in host cell multiplying.

Unattached and attached bacteria and the T-helper lymphocytes, memory B-cells, and lymphocytes are gov-erned by the following differential equations:

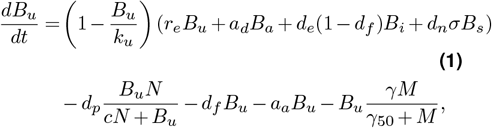

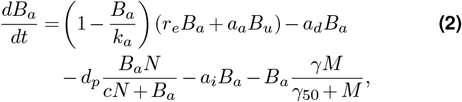

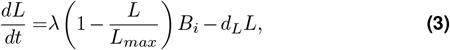

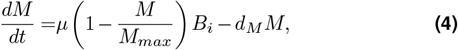

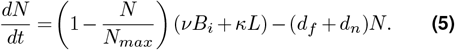

Intracellular gonococcal populations (*B*_*i*_ and *B*_*s*_) are modelled according to the occupancy frequencies of the host cells. WIGWAM simulates the number of host cells containing 0, 1, 2, … bacteria. Host cells transition between groups when bacteria invade or are engulfed, already internalised bacteria replicate or die, or host cells die and are replaced. The governing equations for epithelial cells are:

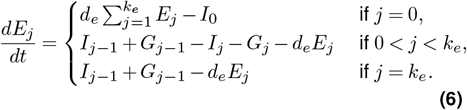

While the total number of epithelial cells is constant, the number of neutrophils varies, depending on recruitment and apoptosis rates. The equations governing neutrophils are:

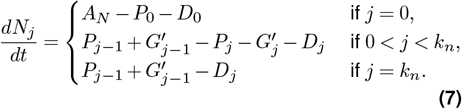

The gonococcal populations in the epithelium, *B*_*i*_ and the gonococcal population surviving within neutrophils, *B*_*s*_ are then:

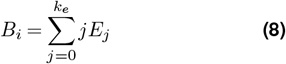

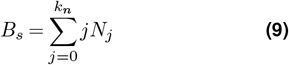

Once the total number of bacteria drops below 10, WIGWAM considers the infection cleared. Section 2 of theflow recruitment and activation killing through IgM phagocytosis epithelial invasion release of intracellular bacteria Supporting Information contains more details on equations S1 and S2, as well as an overview of all variable symbols in equations 1-9 and the ranges of values used for each model parameter (section 4).

WIGWAM simulates the effects of different opacity-associated (Opa) proteins on gonococcal interactions with the epithelium and neutrophils, see the Supporting Information, section 3. WIGWAM also allows for the specification of different *N. gonorrhoeae* variants, each with their own antimicrobial susceptibility and associated fitness costs. Transitions between different variants happen based on specified mutation rates.

The effect of antibiotics on gonococcal populations is modelled using a Hill equation Goutelle et al. (2008), according to which the effect *E* of the ciprofloxacin concentration *C* at the site of infection (the urinary tract) is equal to

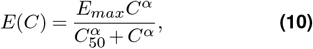

where *E*_*max*_ is the maximum effect, *C*_50_ is the half concentration, at which 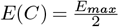, and *α* is the Hill coefficient of sigmoidicity. Ciprofloxacin is a bactericidal antibiotic, so *E*(*C*)*≥* 0 is the kill rate and *−E*(*C*)*B* is added to the equation governing the bacterial population *B* (Equations 1, 2, S1, S2. See also SI section 6). To simulate the effects of varying exposures, including low-level antibiotic exposure, WIGWAM mechanistically describes the absorption, distribution, metabolism, and excretion of the relevant antibiotic within the host, using a modified version of the whole body physiologically-based pharmacokinetic PBPK model described in Sadiq et al. (2017) (see the Supporting Information, section 5). We assume that all bacterial populations are subjected to the ciprofloxacin concentration in the urinary tract, as predicted by this PBPK model.

WIGWAM was parameterised using a Latin hypercube design, with initial parameter ranges based on *in vitro*generated data as described in the literature. All parameter sets that resulted in realistic infection profiles were retained, i.e., untreated infection clearance times between 14 and 200 days Johnson et al. (2010), and daily excreted colony forming units (CFUs) between 9.46 10^4^ and 3.77 10^9^. The latter was based on measured CFU counts van der Veer et al. (2020) and an assumed dailty urine production of 1.5L. Calibration of the PBPK-model was based on published pharmacokinetic data (Figure S1).

### Case study: Low-level ciprofloxacin exposure

We simulated the impact of background exposure to ciprofloxacin on resistance in a population of MSM (men having sex with men). Sequences of 1,000 untreated infections were simulated, in which the dynamics of each infection determined the starting conditions of the next. Transmission happens 1-30 days after the start of each infection, based on the number of recent sexual contacts reported by visitors to an Amsterdam STD clinic. Each infection starts with a random sample of 5% of the bacteria excreted by the infecting partner on the day of infection, with a minimum of 1, 000 bacteria.

We assumed that exposure to ciprofloxacin happens through diet, and daily exposure is divided over three set times each day. Exposure amounts were based on recent studies in China where one study found up to 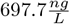 of ciprofloxacin in drinking water in Guangzhou Wang et al. (2010). In Shanghai, up to 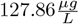 ciprofloxacin was found in urine and a daily ingestion rate of up to 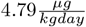 was estimated Wang et al.(2018). Assuming a weight of 77*kg*, this equates to exposures up to 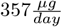.

Ciprofloxacin resistance in *N. gonorrhoeae* is mostly driven by certain single nucleotide polymorphisms (SNPs) in the *gyrA* and *parC* genes Peterson et al. (2015). We assume that each successive level of resistance requires an additional SNP. We use pharmacodynamic properties of different strains to represent successive variants emerging during infection. These strains are, with decreasing susceptibility, DG666 (wildtype), WHO G, WHO M, WHO N, WHO L, and WHO K.Table 1 contains their pharmacodynamic properties Foerster et al. (2016); Unemo et al. (2016). Since we are simulating variants of a single strain, we used the average Hill coefficient of 1.66 for all variants in the model.Bacterial populations transition between subpopulations, each representing a different variant. In order to make the emergence of resistance during an infection stochastic rather than deterministic, we did not allow for populations smaller than a single bacterium. At the end of each timestep, any population smaller than one is set to either zero or one, with a probability according to its value (e.g. 0.25 is set to one with a 25% probability).

**Table 1.** Relevant features of strains used for the parametrisation of ciprofloxacin resistance for simulated variants, listed in order of decreasing susceptibility. The half concentration was calculated from the other parameters, see the supplementary information section 6.

| Strain | DG666 | WHO G | WHO M | WHO N | WHO L | WHO K |
| --- | --- | --- | --- | --- | --- | --- |
| NG-MAST ST | - | ST621 | ST3304 | ST556 | ST1422 | ST1424 |
| zMIC (mg/L) Foerster et al. (2016) | 0.0018 | 0.033 | 0.21 | 1.7 | 6.4 | 18 |
| MIC (mg/L) Unemo et al. (2016) | <0.004 | 0.125 | 2 | 4 | >32 | >32 |
| Resistance rating Unemo et al. (2016) | S | LLR | R | R | HLR | HLR |
| Half concentration ($\frac{mg}{l}$) | 0.012 | 0.058 | 0.313 | 1.453 | 6.895 | 18.164 |
| Max effect (day<sup>-1</sup>) Foerster et al. (2016) | 259.92 | 73.92 | 43.2 | 33.84 | 45.84 | 35.76 |

The mutation rate for *N. gonorrhoeae* was estimated at 8.6. 10^*−*6^ mutations (per site) per year Ezewudo et al. (2015). This translates into mutation rates per site per division of 9.8 10*−*10 in WIGWAM, where non-internalised gonococci divide approximately 24 times per day. This is similar to their doubling time in liquid media. Slightly rounding up the higher of these estimates, we used a mutation rate of 1 · 10^*−*9^ per site per division. To also account for antibiotic-induced increases to the mutation rate, we included higher mutation rates in the simulated scenarios as well. In WIGWAM, resistance mutations are acquired randomly, independent of antibiotic stress, and selection happens solely through differential growth across variants, caused by pharmacodynamics and resistance-associated fitness costs. We assumed resistance mutations do not affect transmission probability. We also assumed mutations back to the wildtype happen with the same probability and revert any associated fitness costs.

### We simulated three scenarios

- Baseline scenario:The first infection is with the wildtype strain, no fitness cost associated with resistance mutations.
- Cost of resistance: The first infection is with the wildtype strain, each consecutive resistance mutation reduces the bacterial growth rate with 2%, relative to the wildtype strain, similar to the growth rate reduction associated with ciprofloxacin resistance observed in other bacterial species Melnyk et al. (2015).
- Resistance present: All resistance mutations are present at the start of the simulation, there is a 2% growth rate reduction per resistance mutation.

For each of these three scenarios, we varied the following parameters:

- Daily ciprofloxacin exposure: 0, 0.015, 0.03, 0.075, 0.15, 0.3,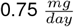.
- Mutation rate: 10^*−*9^, 10^*−*8^, 10^*−*7^ per generation.

For each combination of scenario and parameters, we simulated at least ten sequences of 1,000 consecutive infections. For scenarios with high variance between samples, we simulated additional sequences until the standard error of the mean (as defined in Supplementary Information section 6) was not reduced substantially anymore. This means that the number of sequences per scenario varies between ten and 26 (Supplementary Information section 6, supplemental figures S2 and S3).

## Results

Upon infection, in the absence of antibiotic exposure, the bacterial population grows logistically towards the carrying capacity (Figure 2A). Lymphocytes, memory B cells, and neutrophils are recruited to, and activated at, the site of infection. A proportion of the phagocytosed bacteria survive and multiply in neutrophils. As the immune response is fully activated, extracellular bacteria start to disappear and, as infected epithelial cells are replaced and neutrophils die, intracellular bacteria slowly disappear as well, clearing the infection. Since we assumed ciprofloxacin exposure at three set times during the day, there is a regular pattern of ciprofloxacin concentrations at the site of infection (Figure 2B). While this low-level exposure to ciprofloxacin does not alter the course of infection substantially (Figure 2C), it can lead to shifts towards lower susceptibility (Figure 2D).

**Figure 2.**
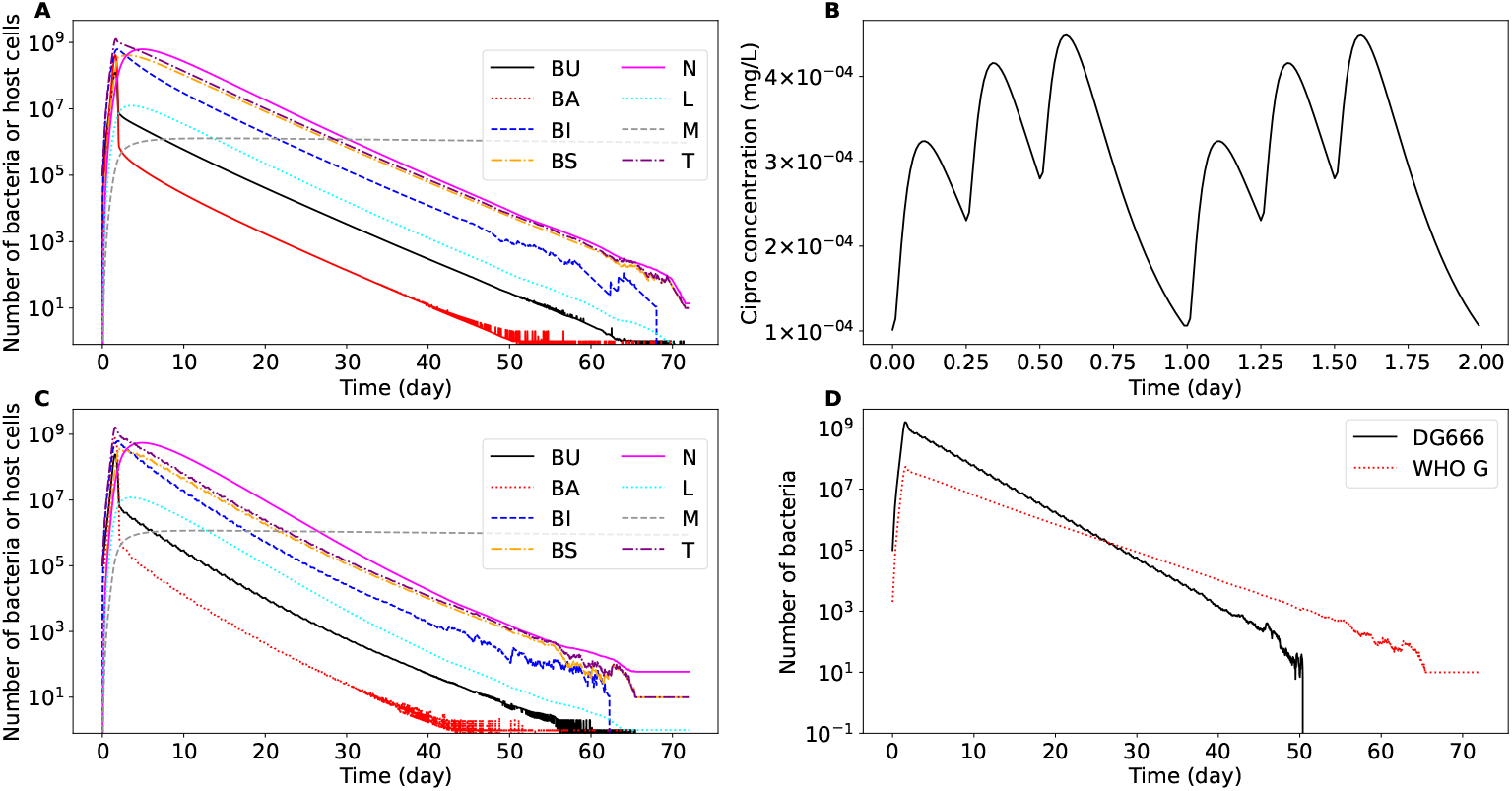
WIGWAM simulations of untreated infections with and without low-level ciprofloxacin exposure. (**A**) A typical untreated infection as simulated by WIGWAM. The lines indicate the sizes of bacterial and host cell populations in the site of infection over time. The oscillations near the end of the simulation are due to stochasticity in the bacterial populations becoming visible at low bacterial populations, this is also the case for (**C**) and (**D**). (**B**) The simulated ciprofloxacin concentration at the site of infection as a result of exposure to 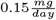 of ciprofloxacin. The ciprofloxacin enters the body at three times during the day (representing three daily meals),leading to a daily varying pattern. (**C**) An infection in an individual exposed to low amounts of ciprofloxacin 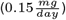. (**D**) The same infection as shown in (**C**) but now the total number of bacteria of two different variants (representing DG666 and WHO G) present are shown. The daily ciprofloxacin exposure drives selection for the less susceptible variant (WHO G) over the wildtype (DG666) variant. BU = unattached bacteria, BA = attached bacteria, BI = bacteria internalised in the epithelium, BS = bacteria surviving inside neutrophils, N = neutrophils, L = lymphocytes, M = memory B cells, T = total bacteria.

### Baseline scenario

During sequences of untreated infections, low-level exposure to ciprofloxacin might lead to the development of low-level resistance (LLR) or resistance (R) (Figure 3). The extent of resistance, and the time until resistance mutations are ubiquitous in the bacterial population, depends on the exposure and the rate of mutation. At a mutation rate of 10^*−*9^ mutations per site per generation (MpSpG), LLR-variants became dominant within 100 consecutive infections (equalling approximately 1,400 days), if their host was exposed to at least 0.15 mg ciprofloxacin per day (Figure 3A). At a daily exposure of 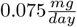, LLR-variants did emerge but did not become dominant. At the highest level of exposure simulated, 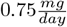, the variant with a second resistance mutation emerged and established in several instances.

**Figure 3.**
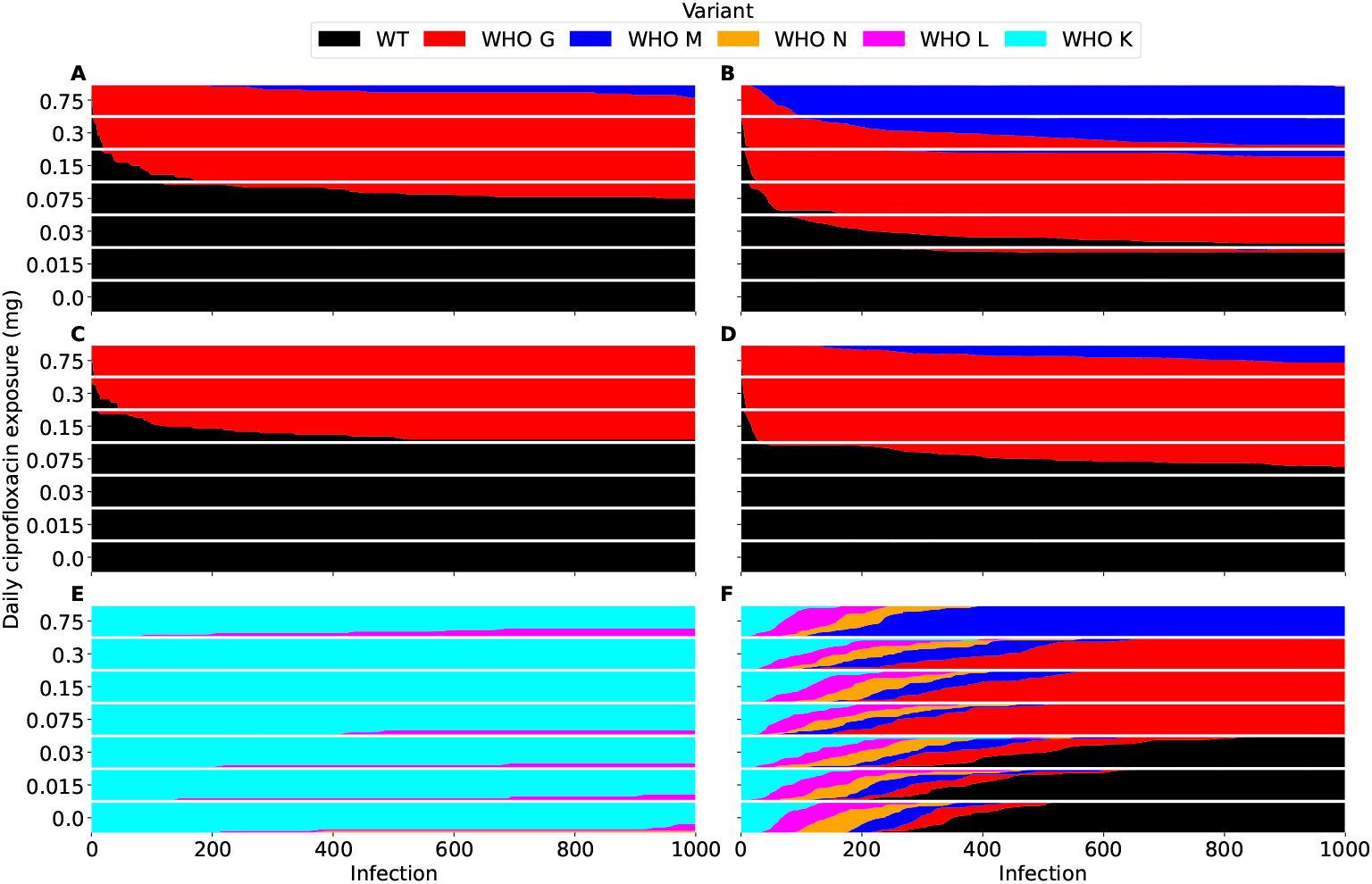
Emergence and selection for resistance during a sequence of 1,000 infections under different daily ciprofloxacin exposures. Each bar shows the distribution of variants across ten or more sequences of 1000 consecutive infections, with the height of each colour indicating the contribution of the corresponding variant. The pharmacodynamic MICs of the variants are 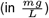 DG666 = 0.0018, WHO G = 0.033, WHO M = 0.21, WHO N = 1.7, WHO L = 6.4, WHO K = 18. Each row of two panels shows the resistance emergence and selection for two different mutation rates (10^*−*9^ (**A, C, E**) and 10^*−*7^ (**B, D, F**)). From top to bottom the three scenarios are: A scenario without cost of resistance, starting with fully susceptible infections (**A, B**), a scenario with a cost of resistance represented by a 2% growth rate reduction per subsequent resistance mutation, starting with fully susceptible infections (**C, D**), and a scenario with the same resistance cost but starting with fully resistant infections (**E, F**).

At a higher rate of 10^*−*7^ MpSpG, ciprofloxacin exposures of 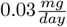already selected for low-level resistance (LLR) (3B). Under a ten-fold higher daily exposure of 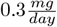, 50% of the bacteria across all repetitions had two or more resistance mutations after 250 infections. At a ciprofloxacin exposure of 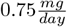, all bacteria were resistant after 100 infections.

### Cost of resistance scenario

If each consecutive resistance mutation reduces the growth rate by 2%, their selective advantage decreases (Figure 3C, 3D). At a rate of 10^*−*9^ MpSpG, 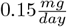 of ciprofloxacin or higher selected for low-level resistance, while high-level resistance did not appear under any of the simulated exposure levels (Figure 3C). At a rate of 10^*−*7^ MpSpG, low-level resistance was already selected for at a daily ciprofloxacin exposure of at least 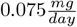 (Figure 3D). Only at the highest simulated exposure of 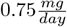, did bacteria with two consecutive resistance mutations emerge and establish.

### Ciprofloxacin resistance prevalent scenario

Assuming a 2% cost of resistance while starting with a bacterial population carrying all resistance mutations, two or less resistance mutations disappeared in each sequence of 1,000 infections, regardless of ciprofloxacin exposure and under a mutation rate of 10^*−*9^ MpSpG, (Figure 3E). While it is beneficial to lose resistance mutations in the absence of ciprofloxacin, the typical growth differential is too small to revert to the susceptible variant. At a rate of 10^*−*7^ MpSpG, there was a reversion to full susceptibility for ciprofloxacin exposures of 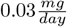and lower (Figure 3F). Exposures from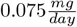 to 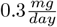 resulted in all low-level resistant variants, while for an exposure of 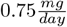 resistance was sustained, in a variant that had lost three resistance mutations. These end states are similar to those of the second scenario (Figures 3D and 3F), indicating that these are equilibrium states where the costs and benefits of resistance cancel each other out.

## Discussion

Unintentional antibiotic exposure can happen via different pathways, such as food, drinking water, and livestock and environmental exposure González et al. (2022); ECDC et al. (2017); Jammoul and El Darra (2019); Ji et al. (2010a,b); Muaz et al. (2018); Shao et al. (2021); Wang et al. (2018); Zhou et al. (2021), and this can lead to bacterial exposure to antibiotics above minimal selective concentrations Subirats et al. (2019).

The ciprofloxacin exposure levels we tested led to concentrations varying from 1.02.10^*−*5^ to 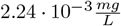at the site of infection. In vitro experiments showed that ciprofloxacin concentrations as low as 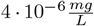 can select for antimicrobial resistance González et al.(2022). This implies that even lower exposures than we simulated potentially select for resistance in a population, providing an explanation for the observed nondecreasing trend of ciprofloxacin resistance in *N. gonorrhoeae* in the absence of treatment Hooshiar et al. (2024). This is despite the fact that bacteria carrying resistance mutations have to be transmitted in order to spread through a population, the probability of which depends on the within-host selection pressure and time between transmissions.

Since ciprofloxacin resistance is associated with fitness costs in several bacterial species Melnyk et al. (2015), it is likely there is a similar fitness cost in *N. gonorrhoeae*. Our simulations show that with a mutation rate of 10^*−*9^ mutations per site per generation, such a fitness cost does not provide enough selection pressure for reversion to the susceptible variant (Figure 3E). Besides background exposure, there are other potential explanations for resistance mutations persisting, such as compensatory mutations negating fitness costs associated with resistance mutations Schulz zur Wiesch et al. (2010) or the fact that certain combinations of resistance mutations can result in a fitness advantage over susceptible strains Helekal et al. (2025). There might also be bystander exposure, the use of ciprofloxacin for treatment of other infections in people with (asymptomatic) gonorrhoea Kenyon et al. (2020); Olesen and Grad (2020). Finally, since antibiotic use by highly sexually active MSM is frequent, co-resistance due to shared mechanisms such as efflux pump upregulation can explain the persistence of ciprofloxacin resistance Blair et al. (2014).

Our results are sensitive to the mutation rate and costs associated with resistance. We assumed the rates at which SNPs occur reported in the literature Ezewudo et al. (2015) apply to the *gyrA* and *parC* genes, which might not be the case. We also made the simplifying assumption that the rate at which resistance mutations appear is the same as the rate at which they are reverted, although we know that mutation rates are not necessarily equal across genomes and base pairs Lind and Andersson (2008). The only ciprofloxacin resistance related cost we took into account was a reduction in growth rate similar to those for *E. coli* and *S. pneumoniae* reported by Melnyk Melnyk et al. (2015), while transmissibility could also be affected. In addition, while occasional exposure to doses as high as 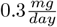 is plausible Wang et al. (2018), daily unintentional exposure to such amounts is unlikely, one would expect high variation in exposure levels, both between people and over time. On the other hand, we simulated a scenario with only one infected person at any given time, passing on the infection to just one other person, which can be considered a conservative approach. Since environmental and dietary sources of antibiotics could expose many people, there are more opportunities for resistance to emerge and be selected for, in addition to social network dynamics such as highly promiscuous people potentially rapidly spreading resistant strains. Finally, longterm, asymptomatic infections are common and make within-host selection of resistance more likely than the short-term infections we considered.

There are some inherent model limitations that should be noted. First of all, for model output validation there is only limited data available in the form of single colony-forming unit measurements in the urine of infected people van der Veer et al. (2020), as opposed to full time series measurements of within-host bacterial populations. There are also still many unknowns regarding gonorrhoea, such as what causes an infection to be symptomatic and the interactions between the bacteria and the immune system, as well as the values of certain parameters, which necessitated making assumptions in the construction of this model.

Rather than relying on epidemiological data, available only for antibiotics already in use, mechanistic withinhost approaches as presented here allow us to study the likelihood of resistance emerging for novel antibiotics and stewardship scenarios by leveraging existing knowledge. The mechanistic nature means that to adapt our model for use with different antibiotics, such as the currently recommended first-line treatment ceftriaxone, one only has to collect the relevant pharmacokinetic and pharmacodynamic parameters and create a model representing the emergence or acquisition of resistance. Besides predicting the emergence of resistance in potential treatment and stewardship scenarios, mechanistic approaches are also able to account for factors such as heterogeneity in the target population or varying transmission probabilities more accurately than SIR models Smieszek (2009). Additionally, as we have demonstrated here, mechanistic models are suitable for studying the emergence of resistance due to unintentional exposure, yielding insights into the risks for different exposure scenarios. As such, mechanistic approaches are a valuable tool in the global health and epidemiology contexts.

Our results highlight the importance of secure disposal of antibiotics, proper water treatment, and food safety regulations in order to avoid unintentional exposure to antibiotics. The One Health approach, which recognises the links between human, animal, and environmental health, has been recognised as an important public health framework, especially where antibiotic resistance is concerned Velazquez-Meza et al. (2022), which is underlined by this study, even for infections with human-host restricted pathogens, such as *N. gonorrhoeae*.

