## Supplemental Information for "Environmental exposure to antibiotics drives and sustains antibiotic resistance in *Neisseria gonorrhoeae*: A modelling study"

### Supplementary Information

#### Supplementary Note 1: Technical details of WIGWAM

WIGWAM is written in C++ and requires input files (in XML-like format) to run. Such an input file specifies simulation parameters such as runtime, timestep, numerical integration method and output printing parameters. The input file also specifies which models should be used for each part of the model (such as the antibiotic kinetics), as well as relevant parameters and general parameters which should be used in the simulation. Model parameters are not hard coded to allow for flexibility. WIGWAM output is printed to text files, in csv or txt format. One file contains diagnostic, warning and error messages, one file contains a record of the model state at specified intervals, one file contains random samples of excreted bacteria which represent populations that sexual contacts would be infected by and one file contains a high level overview of the infection dynamics at specified intervals. Each output can be turned on or off.

Each state variable in a WIGWAM simulation, representing model elements such as the time, the number of unattached bacteria or the antibiotic concentration at the site of infection, is governed by either a differential equation, an algebraic equation or is given as an input parameter. The state of the simulation is stored in a state vector $S$, the first entry of which is the time $t$. WIGWAM allows the use of different numerical integration methods to integrate differential equations forward in time. The recommended method, used in this paper, is the adaptive Runge-Kutta-Fehlberg method, an adaptive method of order $O(h^4)$ with an error estimator of order $O(h^5)$. The numerical integration method is used to calculate the state of the model at time $t+\Delta t$, with algebraic expressions evaluated at each step of the numerical integration. The code associated with each state variable uses the state vector to calculate the rate of change of that state variable (or the value of that state variable in case its governed by an algebraic equation). Output is written at the specified time intervals, with values linearly interpolated in case the timestepping of the numerical integration method does not line up with the output writing times.

#### Supplementary Note 2: Equations governing intracellular bacterial populations

The differential equations governing the evolution of intracellular bacterial populations are

$$\frac{dE_j}{dt} = \begin{cases} d_e \sum_{j=1}^{k_e} E_j - I_0 & \text{if } j=0, \\ I_{j-1} + G_{j-1} - I_j - G_j - d_e E_j & \text{if } 0<j<k_e, \\ I_{j-1} + G_{j-1} - d_e E_j & \text{if } j = k_e. \end{cases} \tag{S1}$$

and

$$\frac{dN_j}{dt} = \begin{cases} A_N - P_0 - D_0 & \text{if } j=0, \\ P_{j-1} + G'_{j-1} - P_j - G'_j - D_j & \text{if } 0<j<k_n, \\ P_{j-1} + G'_{j-1} - D_j & \text{if } j = k_n. \end{cases} \tag{S2}$$

Here $I_n$ is the rate at which bacteria invade epithelial cells containing $n$ bacteria, $G_n$ is the bacterial growth rate for epithelial cells containing $n$ bacteria, $A_N$ is the neutrophil activation rate, $P_n$ is the phagocytosis rate of neutrophils containing $n$ bacteria, $D_n$ is the rate at which neutrophils containing $n$ bacteria disappear (through being excreted from the body or apoptosis), and $G'_n$ is the bacterial growth rate for neutrophils containing $n$ bacteria. When written out in full, these equation are:

$$\frac{dE_j}{dt} = \begin{cases} d_e \sum_{j=1}^{k_e} E_j - a_i \frac{E_0}{n_e} B_a & \text{if } j=0, \\ a_i B_a \frac{E_{j-1}}{n_e} + r_i E_{j-1}(j-1)\left(1-\frac{j-1}{k_e}\right) f^{log}_{k_e,j-1} - a_i B_a \frac{E_j}{n_e} - r_i E_j j \left(1-\frac{j}{k_e}\right) f^{log}_{k_e,j} - d_e E_j & \text{if } 0<j<k_e, \\ a_i B_a \frac{E_{j-1}}{n_e} + r_i E_{j-1}(j-1)\left(1-\frac{j-1}{k_e}\right) f^{log}_{k_e,j-1} - d_e E_j & \text{if } j = k_e. \end{cases} \tag{S3}$$

Here $E_j$ is the group of epithelial cells containing $j$ bacteria, $t$ is the time, $d_e$ is the epithelial cell replacement rate, $k_e$ is the maximum number of bacteria per epithelial cell, $a_i$ is the rate at which attached bacteria enter the epithelial cells, $n_e$ is the number of epithelial cells, $B_a$ is the population of attached bacteria, $r_i$ is the growth rate of bacteria in epithelial cells and $f^{log}_{k_e,j}$ is a correction factor for the growth rate. The equation for the neutrophils is:

$$\frac{dN_j}{dt} = \begin{cases} \left(1-\frac{N_{tot}}{N_{max}}\right)(\nu B_i+\kappa L) - d_p s N_0 \frac{B_u+B_a}{cN_{tot}+B_u+B_a} - (d_f+d_N)N_0 & \text{if } j=0, \\ d_p s(N_{j-1}-N_j)\frac{B_u+B_a}{cN_{tot}+B_u+B_a} + r_s N_{j-1}(j-1)\frac{k_n-j+1}{k_n} f^{log}_{k_n,j-1} & \\ -r_s N_j j \frac{k_n-j}{k_n} f^{log}_{k_n,j} - (d_f+d_N)N_j & \text{if } 0<j<k_n, \\ d_p s N_{j-1}\frac{B_u+B_a}{cN_{tot}+B_u+B_a} + r_s N_{j-1}(j-1)\left(1-\frac{j-1}{k_n}\right) f^{log}_{k_n,j-1} - (d_f+d_N)N_j & \text{if } j=k_n. \end{cases} \quad \textbf{(S4)}$$

Here $N_j$ is the group of neutrophils containing $j$ bacteria, $N_{tot}$ is the total number of neutrophils, $N_{max}$ is the maximum number of neutrophils, $\nu$ is the rate at which bacteria in epithelial cells lead to the recruitment of neutrophils, $B_i$ is the population of bacteria in epithelial cells, $\kappa$ is the rate at which lymphocytes recruit neutrophils, $L$ is the population of lymphocytes, $d_p$ is the rate at which neutrophils phagocytose bacteria, $s$ is the fraction of phagocytosed bacteria that survives in neutrophils, $B_u$ is the population of unattached bacteria, $c$ is a constant, $d_f$ is the rate at which neutrophils are excreted from the urethra, $d_N$ is the rate at which neutrophils die, $r_s$ is the growth rate of bacteria in neutrophils and $f^{log}_{k_n,j}$ is a correction factor for the growth rate.

These equations contain the term $f^{log}_{k_e,j}$, which accounts for the fact that these equations describe the transition rates of cells containing a certain number of bacteria, instead of the growth rate of the bacterial population. $f^{log}_{k_e,j}$ is equal to:

$$f^{log}_{k_e,j} = -\frac{1}{\ln(\frac{k_e j - j - j^2}{k_e j + k_e - j - j^2})}. \quad \textbf{(S5)}$$

We model the growth of a population of intracellular bacteria $P$ with the logistic growth equation

$$\frac{dP}{dt} = rP(1-\frac{P}{k}). \quad \textbf{(S6)}$$

where $r$ is the maximum growth rate and $k$ is the carrying capacity. This has the analytic solution

$$P(t) = \frac{kP_0}{P_0+(k-P_0)e^{-rt}}. \quad \textbf{(S7)}$$

With this expression for $P(t)$, we set $P_0 = n$ and $P(t) = n+1$ to get

$$n+1 = \frac{kn}{n+(k-n)e^{-rt}} \Rightarrow n+(k-n)e^{-rt} = \frac{kn}{n+1} \Rightarrow e^{-rt} = \frac{1}{k-n}(\frac{kn}{n+1}-n) \quad \textbf{(S8)}$$

$$\Rightarrow e^{-rt} = \frac{1}{k-n}\frac{kn-n-n^2}{n+1} \Rightarrow e^{-rt} = \frac{kn-n-n^2}{kn+k-n^2-n} \quad \textbf{(S9)}$$

$$\Rightarrow t = -\frac{\ln(\frac{kn-n-n^2}{kn+k-n-n^2})}{r}. \quad \textbf{(S10)}$$

Giving us the growth rate multiplier

$$f^{log}_{k_e,j} = -\frac{1}{\ln(\frac{k_e j - j - j^2}{k_e j + k_e - j - j^2})}. \quad \textbf{(S11)}$$

Note that when $j = 1$, we set $f^{log}_{k_e,j} = 1$.

#### Supplementary Note 3: Opa proteins

The opacity-associated (Opa) proteins that gonococci present on their outer membrane, undergo rapid phase variation. These Opa proteins mediate intercellular reactions between gonococci and human host cells, including epithelial cell invasion and phagocytosis by neutrophils. Each of the 11 Opa independently undergoes phase variation Bhat et al. (1991); Dempsey et al. (1991) with an estimated frequency of $\phi = 10^{-3} - 10^{-4}$cell$^{-1}$generation$^{-1}$ *in* *vitro* Mayer (1982); Murphy et al. (1989). At any time, a gonococcal infection thus consists of a mixture of bacteria expressing one, none, or multiple distinct Opa proteins.

Considering that gonococci must express Opa proteins to allow for tight interaction and epithelial cell invasion Kupsch et al. (1993), we assumed that Opa-negative (Opa-) gonococci, i.e., those that do not express any Opa proteins, do not invade epithelial cells and remain extracellular. In addition, the lack of tight binding makes them susceptible for exfoliation and detachment back into the unattached state Muenzner et al. (2005, 2010), as reflected by differentiation in the detachment rate of attached NG, based on the Opa expression. Indeed, a strong selection pressure for expressing opa genes exists in vivo, as infections with Opa- gonococci in short time lead to Opa-positive (Opa+) gonococci in urine and urethral swabs of male volunteers Jerse et al. (1994). Simultaneously, Opa proteins also mediate the internalization of gonococci in neutrophils and subsequent phagocytosis. As such, gonococci expressing Opa proteins survive less well in the presence of neutrophils in vitro than those that do not express them Criss et al. (2009); Fischer and Rest (1988); Rest et al. (1982). Thus, it has been suggested that the phase variation of individual Opa-encoding genes allows *N. gonorrhoeae* to partially evade rapid killing by neutrophils and sustain infection longer Criss and Seifert (2012); Criss et al. (2021), while at the same time still allowing the exploitation of the intracellular epithelial space for colonization.

The majority of Opa proteins are recognized by one or more human carcinoembryonic antigen-related cell adhesion molecule (CEACAM) receptors Gray-Owen and Blumberg (2006); Sadarangani et al. (2011)(Gray-Owen & Blumberg, 2006; Sadarangani et al., 2011). Of this CEACAM family of receptors, one variant – CEACAM3 – is expressed by neutrophils but not by epithelial cells. Despite the co-presence of other CEACAM variants on neutrophils, CEACAM3 has been shown essential for neutrophils to effectively phagocytose CEACAM-binding, Opa-expressing, bacteria Chen and Gotschlich (1996); Pils et al. (2008)(Chen & Gotschlich, 1996; Pils et al., 2008). Interestingly, a subset of Opa proteins are recognized by other CEACAMs but not by CEACAM3 Bos et al. (1997); Gray-Owen et al. (1997)(Bos et al., 1997; Gray-Owen et al., 1997). By infecting transfected HeLa cell lines expressing individual CEACAM receptors, with recombinant *N. gonorrhoeae* expressing defined opa genes, Gray-Owen et al. (1997)Gray-Owen et al. (1997) showed that 6 out of 11 Opa variants do not interact with CEACAM3 receptors, while all 11 Opa variants bind to the CEACAM variants expressed on epithelial cells. Consequently, we hypothesize that gonococci expressing one or more of these six Opa proteins, but no others, should be capable of epithelial cell invasion while partially avoiding neutrophil recognition and subsequent phagocytosis. Thus, the rate at which a bacterium is phagocytosed depends on the combination of Opa proteins it expresses.

As opposed to tracking the expression of each Opa protein (which requires $2^{11}$ unique states), we group gonoccoci into three behaviourally distinct groups, based on their Opa expression states: bacteria not expressing any Opa proteins, bacteria expressing at least one Opa protein but none that bind to CEACAM3, and bacteria expressing at least one CEACAM3-binding Opa protein. We subdivide each bacterial population (Bu, Ba, Bi, Bs) into these three functionally distinct groups, and use the phase variation rate $\phi$ and the transition matrix $P$, as described in the supplemental information, to determine per bacterial population the transition rates between these three groups . To this end, a transition term is added to each differential equation governing a bacterial population. For the population $B_u^p$, where the superscript $^p$ indicates the OPA state, this term would be:

$$\sum_q (\phi_{q,p} B_u^q - \phi_{p,q} B_u^p) \tag{S12}$$

We introduce the notation $(n1, n2)$ to represent the state where $n1$ Opa proteins which attach to the CEACAM3 receptor are expressed and $n2$ Opa proteins which do not attach to the CEACAM3 receptor are expressed. With this simplification, not taking into account which specific Opa proteins are expressed we represent the $2^{11} = 2048$ unique Opa states in $6 \cdot 7 = 42$ Per generation, the probability to transition from a start state $(n1_s, n2_s)$ to an end state $(n1_e, n2_e)$ is equal to:

$$P((n1_s, n2_s) \to (n1_e, n2_e)) = P(n1_s \to n1_e) * P(n2_s \to n2_e). \tag{S13}$$

Since the probabilities are independent. Then, defining $n1_d = n1_s - n1_e$,

$$P(n1_s \to n1_e) = \sum_{i=0}^{\lfloor \frac{n1 - |n1_d|}{2} \rfloor} P_t(|n1_d| + 2 * i) N(n1_s, n1_e, |n1_d| + 2i). \tag{S14}$$

Here $P_t(n)$ is the probability of $n$ transitions happening in a single generation, which is

$$P_t(n) = f_0^n (1 - f_0)^{n1-n}. \tag{S15}$$

And $N(s, e, n)$ is the number of possible ways to transition from $s$ to $e$ Opa proteins being expressed with a total of $n$ Opa protein expressions switching state, which is

$$N(n_s, n_e, |n_d| + 2i) = \begin{cases} \binom{n_s}{n_d}\binom{n_e}{i}\binom{n-n_s}{i} & \text{if } n_s > n_e \\ \binom{n-n_s}{n_d}\binom{n_s}{i}\binom{n-n_e}{i} & \text{if } n_s < n_e. \end{cases} \tag{S16}$$

From this we can calculate the transition matrix $P$, the entries of which encode the transition probabilities into and out of each state. These matrix $P$ encodes the rate of change per generation for all the different Opa protein states. For a population $A$ with growth rate $r$ we have

$$\frac{dA}{dt} = rA, \tag{S17}$$

which has the solution

$$A = e^{rt} A. \tag{S18}$$

The doubling time for this population is given by

$$e^{rt} = 2 \Rightarrow t = \frac{\log(2)}{r}, \tag{S19}$$

so the number of generations per unit of time $\gamma$ is given by

$$\gamma = \frac{r}{\log(2)}. \tag{S20}$$

This yields

$$\frac{dA}{dt} = P \frac{r(t)}{\log(2)} A. \tag{S21}$$

Using the superscript $^p$ to indicate the Opa protein state and $^{tot}$ to indicate the sum over all Opa states, this means that the equations for the extracellular bacterial populations are

$$\frac{dB_u^p}{dt} = \left(1 - \frac{B_u^{tot}}{k_u}\right)\left(r_e \cdot B_u^p + a_d^p \cdot B_a^p + \sigma_e \cdot d_e(1-d_f)B_i^p + \sigma_n \cdot d_N \cdot B_s^p\right) \tag{S22}$$

$$- d_p^p \frac{B_u^p \cdot N}{c \cdot N + B_u^p} - d_f \cdot B_u^p - a_a \cdot B_u^p - \gamma \cdot M \cdot B_u^p + \sum_q (\phi_{q,p} B_u^q - \phi_{p,q} B_u^p) \tag{S23}$$

$$\frac{dB_a^p}{dt} = \left(1 - \frac{B_a^{tot}}{k_2}\right)\left(r_e \cdot B_a^p + a_a \cdot B_u^p\right) - a_d^p \cdot B_a^p - d_p^p \frac{B_a^p \cdot N}{c \cdot N + B_a^p} - a_i^p \cdot B_a^p - \gamma \cdot M \cdot B_a^p + \sum_q (\phi_{q,p} B_a^q - \phi_{p,q} B_a^p) \tag{S24}$$

The equations governing the intracellular bacterial populations are altered similarly, with an Opa protein transition term.

### Supplementary Note 4: Parameter ranges

Table 1 lists the symbols used for model parameters, as well as a short description of the meaning of each parameter, the unit and the range we assume the parameter to have. These ranges are based on literature values where possible, as explained in this section, with the rest of the parameter ranges taken to be several orders of magnitude in order to maximise the likelihood of containing the right value.

| Symbol | Parameter description | Range | Unit | References |
|---|---|---|---|---|
| $a_a$ | NG attachment rate to epithelial cells | $4.32 - 8.64$ | $day^{-1}$ | Gubish Jr et al. (1979) |
| $a_{d+}$ | Detachment rate of NG with OPA proteins | $0.0288 - 0.264$ | $day^{-1}$ | Muenzner et al. (2005) |
| $a_{d-}$ | Detachment rate of NG without OPA proteins | $1.008 - 1.104$ | $day^{-1}$ | Muenzner et al. (2005) |
| $a_i$ | Rate of NG entering epithelium | $21.6 - 45.36$ | $day^{-1}$ | Shaw and Falkow (1988) |
| $c$ | Ratio dependent constant (used because phagocytosis is modelled as a ratio-dependent predator-prey interaction, see Jayasundara et al. (2019)) | $0.76 - 1.03$ | $b \cdot n^{-1}$ | |
| $d_e$ | Epithelial cell replacement rate | $\frac{ln(2)}{14} - ln(2)$ | $day^{-1}$ | - |
| $d_f$ | Flushing rate | $0.01 - 0.25$ | $day^{-1}$ | - |
| $d_L$ | Lymphocyte/macrophage death rate | $\frac{ln(2)}{140} - \frac{ln(2)}{1.4}$ | $day^{-1}$ | Jones et al. (2015) |
| $d_M$ | Memory B cell death rate | $\frac{ln(2)}{140} - \frac{ln(2)}{1.4}$ | $day^{-1}$ | Jones et al. (2015) |
| $d_N$ | Neutrophil death rate | $\frac{2}{3}ln(2) - 4ln(2)$ | $day^{-1}$ | Liew and Kubes (2019); Simons et al. (2006); Pillay et al. (2010); Colotta et al. (1992); Kim et al. (2011); Simons et al. (2006) |
| $d_p$ | Neutrophil phagocytosis rate | $62.88 - 96.24$ | $\frac{b}{day \cdot n}$ | |
| $d_{pm}$ | Neutrophil phagocytosis rate modifier for NG without CEACAM3 OPA proteins | $0 - 1$ | $-$ | - |
| $k_e$ | Carrying capacity of epithelial cells | $10 - 30$ | $b$ | Veale et al. (1979) |
| $k_{eb}$ | Number of bacteria which can attach externally to epithelial cells | $10 - 300$ | $b$ | Gubish Jr et al. (1979); Mårdh and Westtöm (1976) |
| $k_n$ | Carrying capacity of neutrophils | $5 - 25$ | $b$ | Veale et al. (1979) |
| $k_u$ | Unattached NG carrying capacity | $7.68 \cdot 10^8 - 7.776 \cdot 10^{12}$ | $b$ | - |
| $L_{max}$ | Max leukocytes/macrophages | $10^7 - 10^9$ | $n$ | Sender et al. (2023) |
| $M_{max}$ | Max memory B cells | $10^7 - 10^9$ | $n$ | Sender et al. (2023) |
| $N_{max}$ | Max neutrophils | $1.1 \cdot 10^{10} - 4.4 \cdot 10^{10}$ | $n$ | Sender et al. (2023) |
| $n_e$ | Number of epithelial cells in urethra | $5.12 \cdot 10^6 - 1.44 \cdot 10^8$ | $n$ | - |
| $r_e$ | Growth rate of extracellular NG | $10.32 - 23.52$ | $day^{-1}$ | Foerster et al. (2016) |
| $r_i$ | Growth rate of NG in epithelium | $6.96 - 11.52$ | $day^{-1}$ | Shaw and Falkow (1988) |
| $r_s$ | Growth rate of NG in neutrophils | $7.45 - 9.77$ | $day^{-1}$ | Simons et al. (2005) |
| $s$ | Fraction of NG surviving phagocytosis by neutrophils | $0.01 - 0.5$ | $-$ | - |
| $\gamma$ | Extracellular NG killing rate by memory B cells | $1 - 10000$ | $\frac{b}{n \cdot day}$ | - |
| $\kappa$ | Macrophage/lymphocyte neutrophil recruitment rate | $1 - 1000$ | $day^{-1}$ | - |
| $\lambda$ | Lymphocyte/macrophage activation rate | $10^{-7} - 10$ | $\frac{n}{b \cdot day}$ | - |
| $\mu$ | Memory B cell activation rate | $10^{-6} - 10$ | $\frac{n}{b \cdot day}$ | - |
| $\nu$ | Neutrophil recruitment rate by epithelium | $10^{-8} - 1$ | $\frac{n}{b \cdot day}$ | - |
| $\sigma$ | Fraction of NG in neutrophils which survive neutrophil apoptosis | $0.25 - 1$ | $-$ | - |
| $\phi$ | OPA phase variation rate | $1 \cdot 10^{-4} - 1 \cdot 10^{-3}$ | $gen^{-1}$ | Mayer (1982); Murphy et al. (1989) |

**Table 1.** Symbols used for the parameters in WIGWAM, short description of the parameter, range and unit. $b$ = bacteria, $n$ = number of cells, $gen$ = number of generations.

- $a_a$: This range was obtained by fitting a simple model to the data in Gubish Jr et al. (1979), taking the lower and higher values in that paper as the boundaries of the range.

- $a_i, r_i$: We used experimental data on gonoccocal invasion of human tissue cells and subsequent growth inside these cells Shaw and Falkow (1988). The following simple model was fitted to the data:

$$\frac{dB_a}{dt} = (r_e - a_i) B_a, \tag{S25}$$

$$\frac{dB_i}{dt} = (a_i B_a + r_i B_i) \left(1 - \frac{B_i}{k_e}\right). \tag{S26}$$

  Here the parameters $r_e$ and $k_e$ were varied over their ranges. This yielded a range of values for $a_i$ and $r_i$, and the $2.5$ and $97.5$ percentiles over these values were used as parameter ranges, $21.6$ to $45.36$ for $a_i$ and $6.96$ to $11.52$ for $r_i$.

- $c, d_p$: Following Jayasundara et al. (2019) (see section 1.2.3 of the supplemental information), we use a simplified model describing the unattached bacteria $B_u$ and the phagocytosed bacteria in neutrophils $B_s$, to obtain these parameters by fitting to the data from Rest et al. (1982). The equations used are:

$$\frac{dB_u}{dt} = r_e B_u - d_p \frac{B_u N}{cN + B_u}, \tag{S27}$$

$$\frac{dB_s}{dt} = \left(s d_p \frac{B_u N}{cN + B_u} + r_s B_s\right) \left(1 - \frac{B_s}{N k_n}\right). \tag{S28}$$

  Note that because the experimental data in Rest et al. (1982) was obtained in an experiment lasting 135 minutes, we assume the number of neutrophils remains constant and the bacterial growth is not constrained by capacity.

- $d_e$: The half life of urethral epithelial cells was not found in the literature. NG infection delays apoptosis Binnicker et al. (2003); Massari et al. (2003). We assume that the epithelial cell half life is between 1 and 14 days, so $d_e = \frac{1}{14}\ln(2)$ to $\ln(2)$.

- $d_f$: We assume between $1$ and $25\%$ of unattached bacteria, unattached epithelial cells and neutrophils get flushed out of the urethra every day.

- $d_L, d_M$: The half life of naive follicular and marginal zone B cells is between 13 and 22 weeks while the decay rates for IgM$^+$ and IgG$^+$ memory B cells were markedly slower Jones et al. (2015). However, we want to model how long the memory B cells keep producing IgM, which we assume is shorter than their lifetime. We let this parameter vary from 0.2 to 20 weeks. We use the same range for T helper cells (lymphocytes). So for $d_L$ and $d_M$ the ranges are $\frac{1}{1.4}\ln(2)$ to $\frac{1}{140}\ln(2)$.

- $d_N$: The circulating half life of neutrophils in vivo is considered to be 6-8 hours Liew and Kubes (2019) or 6-12 hours Simons et al. (2006), though one study (whose methodology was criticised) found a half life of 3.75 days Pillay et al. (2010). It was found that during inflammation the half life of neutrophils was found to increased by a bit more than a factor 3 Colotta et al. (1992); Kim et al. (2011). Additionally, *N. gonorrhoeae* delays apoptosis of neutrophils by 12 hours Simons et al. (2006), except when there are a lot of NG in the neutrophil. So we choose as lower and upper limits for the half life 6 and 36 hours (which gives $e^{-1.5d_n} = \frac{1}{2} \Rightarrow d_n = \frac{2}{3}\ln(2)$ to $4\ln(2)$ with a time unit of days).

- $d_{pm}$: It was observed in Criss et al. (2009) that *N. gonorrhoeae* bacteria which did not express CEACAM-binding Opa proteins had higher rates of survival in the presence of neutrophils but the exact reduction in phagocytosis is unclear so we set the range of reduction in phagocytosis as $0$ to $1$.

- $k_e$: Each epithelial cell has their own carrying capacity $k_e$ for internalised bacteria. Experimentally, each infected epithelial cell contained $21.5 \pm 1.7$ NG Veale et al. (1979), so we use $k_e = 10 - 30$.

- $k_{eb}$ bacteria can attach to each epithelial cell externally. In Gubish Jr et al. (1979) the maximum attachment of NG to human cervical cancer cells (HeLa) was about 25 NG per cell. According to Mårdh and Westtöm (1976), the range of number of NG attached to vaginal epithelial cells is $0 - 256$ (with a maximum at a pH of 4.5). So we use $10$ to $300$ as the range.

- $k_n$: Each neutrophil has a carrying capacity of $k_n$. Each infected neutrophil contained $14.7 \pm 1.1$ NG but some contained as many as 50 to $> 100$ Veale et al. (1979). We assume these outliers are not important for the overall dynamics so use $k_n = 10 - 25$.

- $k_u, n_e$: The length, $l_u$ and radius $\rho_u$ of the urethra are $l_u = 2 \cdot 10^5 \mu m$, and $r_u = 8 - 9 \cdot 10^3 \mu m$. The radius of a *N. gonorrhoeae* bacterium is $\rho_g = 0.3 - 0.5 \mu m$ Unemo et al. (2019). Epithelial cells have a radius, $\rho_e$, of $7.5 - 15 \mu m$ as listed here. The diameter of vaginal-cervical cells is $47 \mu m$ (40-50) according to Pearce et al. (1978), so that would imply a radius of $25 \mu m$. So we use $\rho_e = 5 - 25 \mu m$. The carrying capacity for unattached NG bacteria, assuming they exist in the volume of the urethra, is $k_u = \frac{\pi l_u \rho_u^2}{\frac{4}{3}\pi \rho_g^3} = \frac{3 l_u \rho_u^2}{4 \rho_g^3}$. With the numbers given above $k_u = 7.68 \cdot 10^{13}$ to $7.776 \cdot 10^{14}$ but this is assuming the urethra is completely filled with bacteria. We will assume that at most $1\%$ of the urethra volume can be bacteria and take $0.001\%$ of the volume as the lower bound which gives us the bounds: $7.68 \cdot 10^8 - 7.776 \cdot 10^{12}$. The number of epithelial cells in the urethra is equal to $n_e = \frac{2\pi l_u \rho_u}{\pi \rho_e^2} = \frac{2 l_u \rho_u}{\rho_e^2}$. This gives $5.12 \cdot 10^6 - 1.44 \cdot 10^8$.

- $L_{max}, M_{max}, N_{max}$: In Sender et al. (2023) the distribution and amounts of immune cells in the body are listed for each organ. The total number of neutrophils in a reference adult male of 73 kg is $7 \cdot 10^{11}$, most of which are in the bone marrow. In the blood there are about $2.2 \cdot 10^{10}$ neutrophils, while in the "other" tissues, there are very little. So we take as range for $N_{max}$ $1.1 - 4.4 \cdot 10^{10}$. Unfortunately the distribution of immune cells is not given for the urethra specifically, nor is it given for internal epithelial tissues (which the urethra belongs to), but the article does mention that the number of macrophages, T cells and B cells in epithelial tissue varies between $10^6$ and $10^7$ cells per gram of tissue. Taking into account that there are many different types of T cells and B cells and that the T and B cells proliferate in response to exposure to pathogens, we take $10^6$ to $10^8$ cells per gram of tissue as range for the max amount memory B cells and macrophages/helper T cells. Using 10g as the weight of the urethra (assumes a epithelial thickness of 3mm, a urethral diameter of 1mm and a length of 200 mm), we get $10^7 - 10^9$ as ranges for $L_{max}$ and $M_{max}$.

- $r_e$: We based the parameterization of $r_e$, the growth rate of extracellular bacteria (so both unattached and attached) on Foerster et al. (2016). They performed a time-kill curve analysis for multiple antibiotics against a range of N. gonorrhoeae reference strains. We used the growth rates for highly susceptible clinical gonococcal strain DG666 in the absence of ciprofloxacin (two independent experiments), to derive a range of $0.43$ to $0.98 hr^{-1}$.

- $r_s$: According to Simons et al. (2005), 30 minutes after phagocytosis, 58.34% $\pm$ 9.26% of ingested gonococci were viable. This does not account for the gonococci killed by oxidative bursts and only 3.27% of the input CFU were recovered so perhaps more were killed by the neutrophils. Fitting data on the number of active gonococci in neutrophils from the same paper to a simple model with a constant (exponential) growth rate we get a growth rate of 7.45-9.77 day$^{-1}$.

- $s, \gamma, \kappa, \lambda, \mu, \nu, \sigma$: These parameters were not found in the literature so we chose large ranges, spanning several orders of magnitude if possible.

#### Supplementary Note 5: Antibiotic pharmacokinetics

WIGWAM explicitly accounts for antibiotic pharmacokinetics, by mechanistically describing the absorption, distribution, metabolism and excretion of the relevant antibiotic within the host. As such, it allows for the simulation of varying exposure patterns and the associated concentration-time curves of the antibiotic of interest at the sites of infection. For this, we implemented a whole body physiologically-based pharmacokinetic (PBPK) model developed for ciprofloxacin Sadiq et al. (2017), which divides the host's body into multiple compartments that each represent an organ (or collection of organs). Concentrations in each of these compartments are governed by differential equations dependent on the antibiotic concentration in the blood flowing through the compartment, the concentration in the relevant compartment, the blood flow and volume of the compartment and the partitioning coefficient of the antibiotic between blood and the tissue (which is the steady-state ratio between blood and compartment concentration). For some compartments, there are also terms related to drug administration or clearance. Note that while this model was developed for ciprofloxacin, it can be used to model other antibiotics by altering the partition coefficients and relevant administration and clearance parameters.

The differential equation determining the concentration in a tissue Pilari and Huisinga (2010); Sadiq et al. (2017), $C_t$ is

$$\frac{dC_t}{dt} = \frac{Q_t}{V_t}\left(C_a - \frac{C_t}{K_t}\right), \tag{S29}$$

where $Q_t$ is the tissue blood flow, $V_t$ is the volume of the tissue, $C_a$ the arterial concentration and $K_t$ is the tissue partition coefficient, which is equal to the steady state ratio $\frac{C_t}{C_a}$. Equation S29 applies to many organs and tissues but slightly different equations are used for several organs/tissues. These equations are:

$$\frac{dC_a}{dt} = \frac{Q_{co}}{V_a}(C_l - C_a), \tag{S30}$$

$$\frac{dC_v}{dt} = \frac{Q_{co}}{V_v}(\Sigma - C_v), \tag{S31}$$

$$\frac{dC_l}{dt} = \frac{Q_{co}}{V_l}\left(C_v - \frac{C_l}{K_l}\right), \tag{S32}$$

$$\frac{dC_h}{dt} = \frac{1}{V_h}\left(Q_h\left(\Sigma_h - \frac{C_h}{K_h}\right) - C_h\frac{M_h f_{up}}{K_h}\right), \tag{S33}$$

$$\frac{dC_k}{dt} = \frac{1}{V_k}\left(Q_k\left(C_a - \frac{C_k}{K_k}\right) - C_a M_{cr} f_{up}(1 + f_{secr})\right). \tag{S34}$$

Here $C_a$ is the arterial concentration, $C_{co}$ the cardiac output, $V_a$ the arterial volume, $C_l$ the lung concentration, $C_v$ the venous concentration, $V_v$ the venous volume, $V_l$ the lung volume, $K_l$ the lung partition coefficient, $C_h$ the liver concentration, $V_h$ the liver volume, $Q_h$ the liver blood flow, $K_h$ the liver partition coefficient, $M_h$ is the liver clearance rate, $f_{up}$ is the unbound plasma fraction, $C_k$ the kidney concentration, $V_k$ the kidney volume, $Q_k$ the kidney blood flow, $K_k$ the kidney partition coefficient, $M_{cr}$ is the creatine clearance rate and $f_{secr}$ is the estimated fraction of renal clearance that is dependent on secretion from blood to renal tubules.

$$\Sigma = \frac{1}{Q_{co}}\sum_{t\in T} Q_t\frac{C_t}{K_t}, \tag{S35}$$

$$\Sigma_h = \frac{1}{Q_h}\sum_{t\in T_h} Q_t\frac{C_t}{K_t}. \tag{S36}$$

Here $T$ is the set of all tissues connecting into the veins, which is adipose, brain, heart, kidneys, liver, muscle, skin, urinary tract and rest. In other words, $\Sigma$ is the weighed concentration of all tissues from which blood flows directly into the veins. Similarly, $T_h$ is the set of tissues from which blood flows to the liver, which is the spleen, gut and arterial hepatic vein. In order to tailor this model for our specific application, simulating the antibiotic concentration at the urogenital site of infection, we have added a "urinary tract" compartment to the model, the site of infection in this study. Note that this does not include the bladder or the urine, just the urethral tissue. Blood flow, organ volume and partition parameters were taken from the literature or scaled from animal studies if no data on humans is available. The urinary tract concentration as predicted by the PBPK model is the concentration used to calculate the effect on all bacterial populations. Tables 2, 3 and 4 list the compartmental volumes, blood flows and partition coefficients used.
For the calculation of urinary tract blood flow, Delp et al. (1998) lists a urethra blood flow of $7\frac{ml}{100g\cdot min}$ for adult rats and a cardiac index of $28.1\frac{ml}{100g\cdot min}$. This means that the urethral blood flow is $0.25$ times the fractional organ weight times the total cardiac output. Using the numbers from Talja et al. (1991) for piglets and a similar calculation yields $0.4$ times the fractional organ weight times the cardiac output.
After fixing the above volumes, blood flows and partition coefficients for all organs, the ciprofloxacin kidney and liver clearance rates, as well as the absorption rate constant and the oral bioavailability were determined. Tables 5, 6 and 7 list literature values for these parameters.
Based on the values in tables 5, 6 and 7, we set the following bounds for these parameters:

- Ciprofloxacin kidney clearance rate: Between 200 and 700 l/day.
- Ciprofloxacin liver clearance rate: Between 25 and 500 l/day.
- Ciprofloxacin absorption rate constant: Between 15 and 140 1/day.
- Ciprofloxacin oral bioavailability: Between 0.65 and 0.85.

| Compartment | Relative volume female | Relative volume male | Reference |
|---|---|---|---|
| Adipose | 0.3275 | 0.1869 | Sadiq et al. (2017) |
| Arteries | 0.017083 | 0.01918 | Sadiq et al. (2017) |
| Brain | 0.0191781 | 0.01918 | Sadiq et al. (2017) |
| Gut | 0.01644 | 0.01644 | Sadiq et al. (2017) |
| Heart | 0.004167 | 0.004521 | Sadiq et al. (2017) |
| Kidneys | 0.0042466 | 0.004247 | Sadiq et al. (2017) |
| Liver | 0.020724 | 0.02466 | Sadiq et al. (2017) |
| Lungs | 0.00643836 | 0.00643836 | Sadiq et al. (2017) |
| Muscle | 0.2916 | 0.3973 | Sadiq et al. (2017) |
| Rest | 0.20546894 | 0.22174764 | - |
| Skin | 0.03246 | 0.03831 | Sadiq et al. (2017) |
| Spleen | 0.00247 | 0.00247 | Sadiq et al. (2017) |
| Urinary tract | 0.000967 | 0.001041 | Beaudouin et al. (2010) |
| Veins | 0.051248 | 0.05753 | Sadiq et al. (2017) |

**Table 2.** PBPK model compartment relative volumes

| Compartment | Blood flow female | Blood flow male | Reference |
|---|---|---|---|
| Adipose | 0.085 | 0.05 | Sadiq et al. (2017) |
| Arterial hepatic vein | 0.065 | 0.065 | Sadiq et al. (2017) |
| Brain | 0.12 | 0.12 | Sadiq et al. (2017) |
| Cardiac output (CO) | $15 \cdot 24 \cdot BW^{0.74}$ | $15 \cdot 24 \cdot BW^{0.74}$ | Sadiq et al. (2017) |
| Gut | 0.16 | 0.15 | Sadiq et al. (2017) |
| Heart | 0.05 | 0.04 | Sadiq et al. (2017) |
| Kidneys | 0.17 | 0.19 | Sadiq et al. (2017) |
| Liver | 0.255 | 0.245 | Sadiq et al. (2017) |
| Muscle | 0.12 | 0.17 | Sadiq et al. (2017) |
| Rest | 0.149709 | 0.1346877 | - |
| Skin | 0.05 | 0.05 | Sadiq et al. (2017) |
| Spleen | 0.03 | 0.03 | Sadiq et al. (2017) |
| Urinary tract | 0.000291 | 0.0003123 | |

**Table 3.** PBPK model compartment blood flows, all values are as fraction of the cardiac output, while the value listed for cardiac output is in $\frac{L}{h}$. $BW$ is the body weight in kg.

| Compartment | Partition coefficient | Reference |
|---|---|---|
| Adipose | 0.413 | Sadiq et al. (2017) |
| Brain | 0.773 | Sadiq et al. (2017) |
| Gut | 3.35 | Sadiq et al. (2017) |
| Heart | 3.67 | Sadiq et al. (2017) |
| Kidneys | 8.09 | Sadiq et al. (2017) |
| Liver | 3.56 | Sadiq et al. (2017) |
| Lungs | 3.32 | Sadiq et al. (2017) |
| Muscle | 0.977 | Sadiq et al. (2017) |
| Rest | 3.86 | Sadiq et al. (2017) |
| Skin | 0.715 | Sadiq et al. (2017) |
| Spleen | 1.95 | Sadiq et al. (2017) |
| Urinary tract | 3.35 | |

**Table 4.** PBPK model ciprofloxacin partition coefficients.

Note that the nonrenal (liver in the model) clearance rate is listed as higher than the bounds we set here in Gasser et al. (1987). However, renal clearance amounts for most of the ciprofloxacin clearance Vance-Bryan et al. (1990), so we took lower feasible bounds. The measured maximum plasma concentrations, areas under the curves and half times in people for a number of different doses and methods of administration are listed in Vance-Bryan et al. (1990) and Schlender et al. (2018). We numerically minimised the differences between model predictions and these data (Figure S1), resulting in the following values:

- Ciprofloxacin kidney clearance rate: 464 l/day.

| Renal clearance | Rate in $\frac{l}{day}$ | Nonrenal clear | Rate in $\frac{l}{day}$ | Serum clearance | Rate in $\frac{l}{day}$ | Reference |
|---|---|---|---|---|---|---|
| $21.1\frac{l}{h}$ | $506.4$ | – | – | – | – | de Vroom et al. (2020) |
| $245 \pm 101\frac{ml}{min}$ | $353 \pm 145$ | $780 \pm 446\frac{ml}{min}$ | $1123 \pm 642$ | – | – | Gasser et al. (1987) |
| $272 \pm 160\frac{ml}{min}$ | $392 \pm 230$ | $901 \pm 699\frac{ml}{min}$ | $1297 \pm 1006$ | – | – | Gasser et al. (1987) |
| $381 \pm 198\frac{ml}{min}$ | $549 \pm 285$ | $246 \pm 60\frac{ml}{min}$ | $354 \pm 86$ | – | – | Webb et al. (1986) |
| $16.4 \pm 3.5\frac{l}{h \cdot 1.73m^2}$ | $394 \pm 84$ | – | – | $26.8 \pm 5.7\frac{l}{h \cdot 1.73m^2}$ | $643 \pm 137$ | Drusano et al. (1987) |
| $300 - 479\frac{ml}{min}$ | $432 - 690$ | – | – | – | – | Al-Omar (2005) |
| $14.6 \pm 4.5\frac{l}{h \cdot 1.73m^2}$ | $350 \pm 108$ | – | – | – | – | Forrest et al. (1988) |

**Table 5.** Ciprofloxacine clearance rates.

| Absorption rate constant | In $day^{-1}$ | Reference |
|---|---|---|
| $0.718h^{-1}$ | $17.23$ | Meagher et al. (2004) |
| $3.633 \pm 2.170h^{-1}$ | $87.19 \pm 52.08$ | Bergan et al. (1986) |
| $3.367 \pm 2.055h^{-1}$ | $80.81 \pm 49.32$ | Bergan et al. (1986) |
| $3.120 \pm 1.666h^{-1}$ | $74.88 \pm 39.98$ | Bergan et al. (1986) |
| $2.433 \pm 1.886h^{-1}$ | $58.39 \pm 45.26$ | Bergan et al. (1986) |
| $1.78 \pm 5.5h^{-1}$ | $42.7 \pm 132$ | Borner et al. (1986) |
| $2.03 \pm 1.07h^{-1}$ | $48.7 \pm 25.7$ | Borner et al. (1986) |
| $2.14 \pm 1.31h^{-1}$ | $51.4 \pm 31.4$ | Borner et al. (1986) |
| $1.48 \pm 0.5h^{-1}$ | $35.5 \pm 12$ | Borner et al. (1986) |
| $1.84 \pm 0.23h^{-1}$ | $44.2 \pm 5.5$ | Borner et al. (1986) |
| $2.25$ | $54$ | Mean |
| $1.90$ | $45.7$ | Source-aggregated mean |

**Table 6.** Ciprofloxacin absorption rate constant.

| Biological availability | Reference |
|---|---|
| $0.7 - 0.8$ | Farmacotherapeutisch Kompas |
| $0.837$ | Bergan et al. (1986) |
| $0.64$ | Borner et al. (1986) |
| $0.69 \pm 0.07$ | Drusano et al. (1987) |
| $0.69$ | Plaisance et al. (1987) |

**Table 7.** Ciprofloxacine biological availabiliy.

- Ciprofloxacin liver clearance rate: 84.3 l/day.
- Ciprofloxacin absorption rate constant: 44.8 1/day.
- Ciprofloxacin oral bioavailability: 0.663.

We compared how model predictions for ciprofloxacin found in urine compared to literature Boy et al. (2004); Brittain et al. (1985); Naber et al. (1999); Wagenlehner et al. (2006) values (Figure S2). There are also urine ciprofloxacin concentrations found in the literature Gasser et al. (1987), however without knowing the amount of urine collected and the period over which this was collected, these measurements cannot be compared to model predictions in a meaningful way. For this reason we only compare the amount of ciprofloxacin excreted through urine.

### Supplementary Note 6: Antibiotic pharmacodynamics

For the antibiotic pharmacodynamics we use the Hill equation relating the effect $E(C)$, to the antibiotic concentration at the site of infection, $C$ Goutelle et al. (2008).

$$E(C) = \frac{E_{max}C^{\alpha}}{C_{50}^{\alpha} + C^{\alpha}}. \tag{S37}$$

Here, $E_{max}$ is the maximum effect, $C_{50}$ is the half concentration and $\alpha$ the Hill coefficient. Values for $E_{max}$ used for different simulated variants were taken from Foerster et al. (2016) and the Hill coefficient was set to the mean of the Hill coefficients found in the same publication.
Since ciprofloxacin is a bactericidal antibiotic, $E$ is a killing term, subtracted from the bacterial growth equations. So equation S22 becomes (extra term is bold for clarity):

$$\frac{dB_u^p}{dt} = \left(1 - \frac{B_u^{tot}}{k_u}\right)(r_e \cdot B_u^p + a_d^p \cdot B_a^p + \sigma_e \cdot d_e(1-d_f)B_i^p + \sigma_n \cdot d_N \cdot B_s^p) \tag{S38}$$

$$- d_p^p \frac{B_u^p \cdot N}{c \cdot N + B_u^p} - d_f \cdot B_u^p - a_a \cdot B_u^p - \gamma \cdot M \cdot B_u^p + \sum_q (\phi_{q,p} B_u^q - \phi_{p,q} B_u^p) - \mathbf{B_u^p E(C)}. \tag{S39}$$

$E(C)$ is added to the other equations describing bacterial populations in the same way. For the internal populations, $E(C)$ is multiplied by the correction factor $\frac{-j}{\log \frac{j-1}{j}}$ to account for the fact that these equations describe the amount of cells containing $j$ bacteria, instead of the amount of bacteria and the effect of a bactericidal antibiotic leads to exponential decay. So to equation S3, we add

$$E_j E(C) \frac{j}{\log \frac{j-1}{j}} - E_{j+1} E(C) \frac{j+1}{\log \frac{j}{j+1}} \tag{S40}$$

in the case where $0 < j < k_e$ (the other cases will have only one of these terms), where the first term is the reduction in $E_j$ due to antibiotic killing and the second term is an increase in $E_j$ due to antibiotic killing causing a transition from $E_{j+1}$ to $E_j$.

Note that Foerster et al. (2016) does not provide half concentrations, but it does provide zMICs, where the zMIC is defined as the minimum antibiotic concentration at which there is no bacterial growth. By definition, it is true that

$$0 = r - E(zMIC) = r - \frac{E_{max} zMIC^\alpha}{C_{50}^\alpha + zMIC^\alpha}. \tag{S41}$$

Here $r$ is the bacterial growth rate. From this it follows that

$$r\,(C_{50}^\alpha + zMIC^\alpha) = E_{max} zMIC^\alpha, \tag{S42}$$

$$C_{50}^\alpha = \frac{E_{max}}{r} zMIC^\alpha - zMIC^\alpha, \tag{S43}$$

$$C_{50} = \sqrt[\alpha]{\frac{E_{max}}{r} zMIC^\alpha - zMIC^\alpha}. \tag{S44}$$

Giving us the value for $C_{50}$.

#### Supplementary Note 7: Number of repetitions in case study

In the sequence of asymptomatic infections in the presence of background ciprofloxacin exposure we simulate, there are several sources of stochasticity, such as resistance mutations, time between infections and the bacteria which are transmitted between people. Because of this, we have simulated each scenario 10 times. To check that our results accurately reflect the expected outcome, we quantify how uncertainty varies with the sample size. First, we define the distance between two chains of $1000$ infections, $K_1$ and $K_2$ as

$$d(K_1, K_2) = \sqrt{\sum_{i=1}^{10} \sum_{j=1}^{6} \left(K_1^{100i,j} - K_2^{100i,j}\right)^2}, \tag{S45}$$

where $K_n^{i,j}$ is the relative fraction of variant $j$ at the $i$-th infection for chain $n$. In other words, for each chain of $1000$ infections we create a vector consisting of the relative fractions of each variant and then define the distance between two chains as the Euclidean distance between these two vectors. Using this measure of distance, we can now define the sample variance $Var$ of a set of chains $K = \{K_1, ..., K_n\}$ as

$$Var(K) = \frac{\sum_{i=1}^{n} d\left((K_i, \overline{K}\right)^2}{n-1}, \tag{S46}$$

where $\overline{K} = \frac{\sum_{i=1}^{n} K_i}{n}$ is the mean chain (which has the mean relative fraction of each variant at each infection). Then the standard error of the mean $\sigma_{\overline{K}}$ is

$$\sigma_{\overline{K}} = \sqrt{\frac{Var(K)}{n}}. \quad \textbf{(S47)}$$

Given 10 chains for each scenario, we calculate the standard error of the mean for each subsample of $n$ chains and average this to obtain the relationship between sample size and standard error of the mean (Figure S4). In order to ensure accurate estimates of the mean evolution over time, we simulate chains for each scenario while each extra chain reduces the standard error of the mean by more than $0.01$. With this criterium for generating extra samples, we ended up with up to $26$ repetitions for some scenarios (Figure S5).

#### Supplementary Note 8: Figures

Figure S1 compares WIGWAM predicted pharmacokinetic values to those reported in the literature, while Figure S2 does this specifically for urine ciprofloxacin measurements. The PBPK ciprofloxacin model was fitted to the data shown in this figure using ordinary least squares. Figure S3 shows the predicted ciprofloxacin concentration in different organs and tissues over time after oral ingestion of 500 mg of ciprofloxacin. Figures S4 and S5 show how the standard error of the mean for our results varies based on the sample size. Figures S6, S7 and S8 show the full results for all simulated scenarios. The selection for variants of reduced susceptibility is shown per infection as well as per day. The per day graphs show data until the last day for which all 10 sequences of infections were still active (since the duration of each infection is picked from a distribution randomly, not each sequence will be the same number of days).

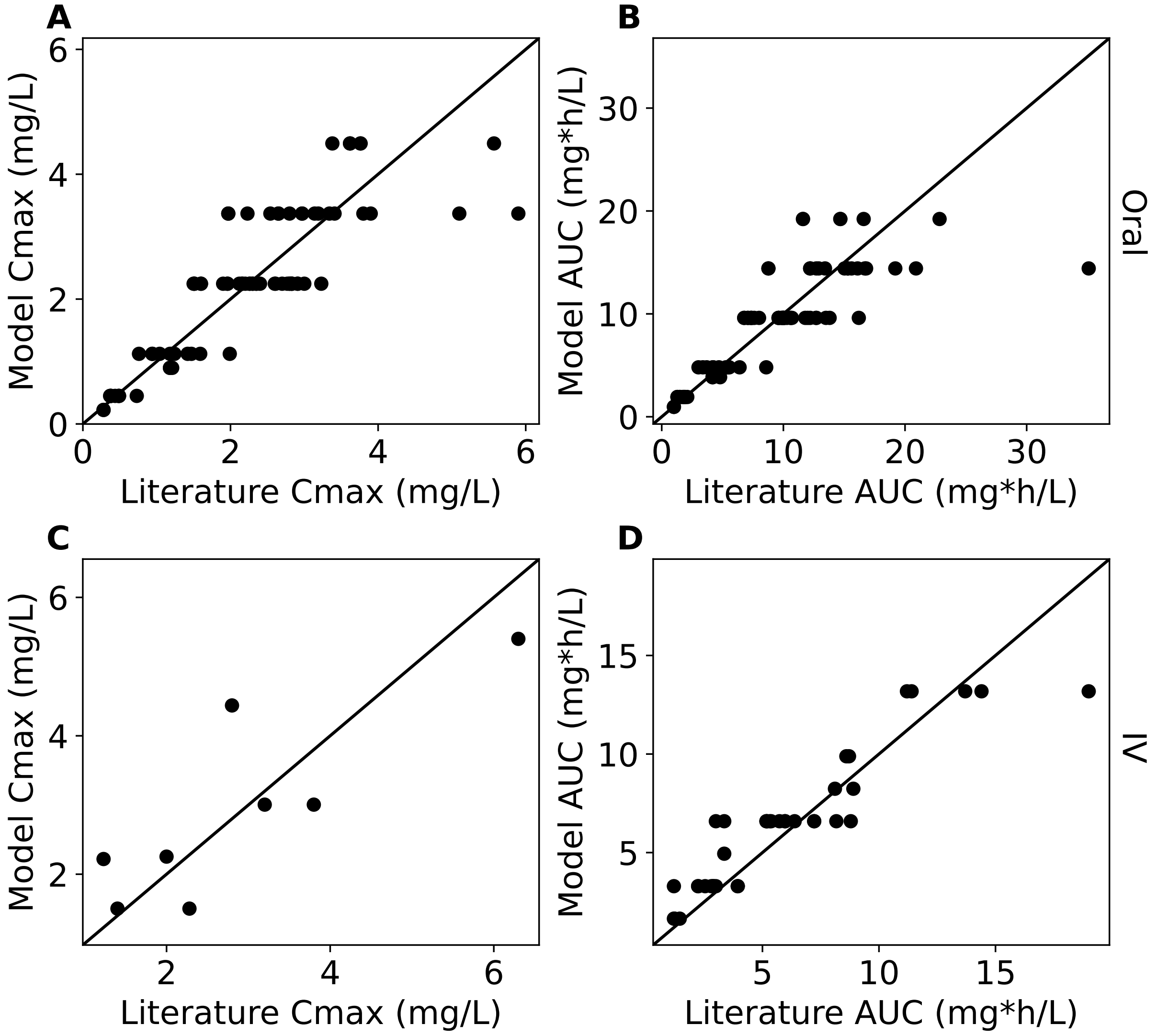


**Figure S1.** Literature pharmacokinetic data compared to the WIGWAM model fitted values based on dose and method of ingestion. The diagonal lines indicate a perfect fit. Panels **A** and **C** compare the maximum antibiotic concentration measured in blood to model fit. Panels **B** and **D** compare the areas under the curve as reported in the literature to model fit. Panels **A** and **B** compare pharmacokinetic values for orally administered ciprofloxacin while panels **C** and **D** compare these values for ciprofloxacin administered intravenously.

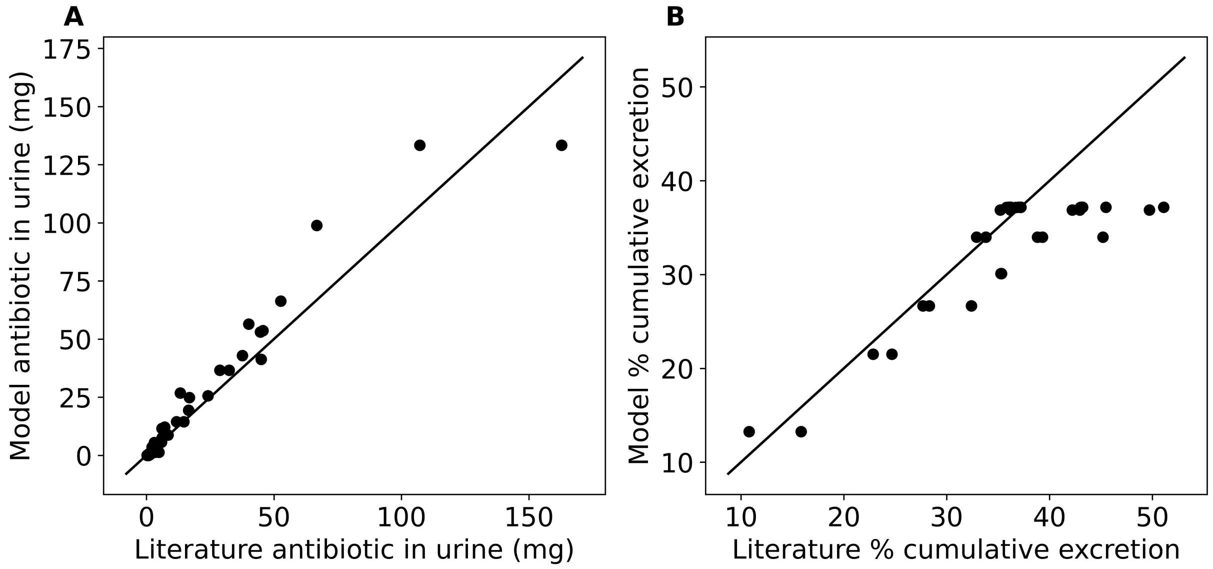


**Figure S2.** Literature urine ciprofloxacin data compared to the WIGWAM model fitted values based on dose and method of ingestion. The diagonal lines indicate a perfect fit. Panel **A** compares the total ciprofloxacin in urine collected over a period of time to model predictions. Panel **B** compares the cumulative percentage of ciprofloxacin excreted in urine to model fit.

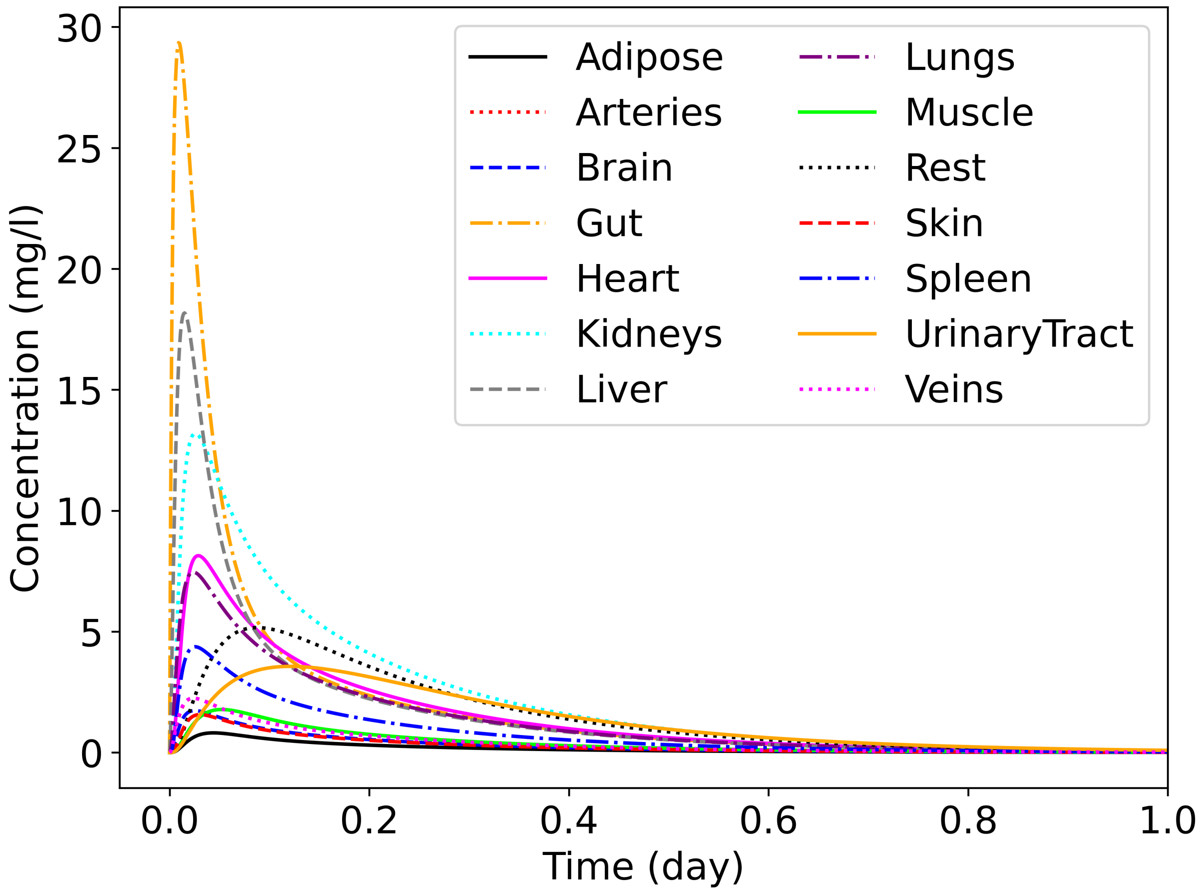


**Figure S3.** Ciprofloxacin concentration in different organs and tissues after oral ingestion of 500 mg of ciprofloxacin as simulated by the WIGWAM PBPK model.

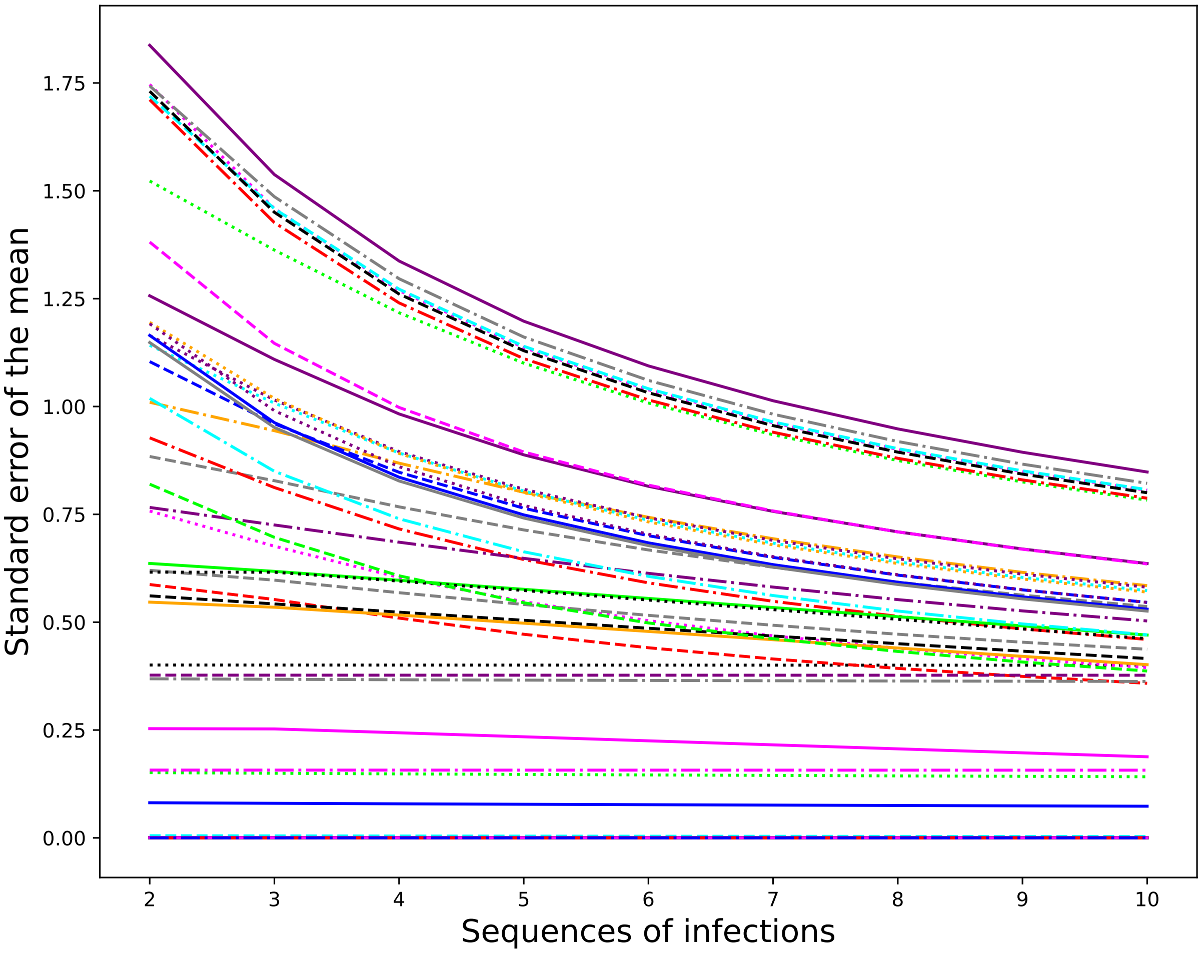


**Figure S4.** The standard error of the mean (as defined in equations S45, S46 and S47) as a function of the sample size. Each line represents a combination of scenario, ciprofloxacin exposure amount and mutation rate. The values are obtained by averaging the standard error of the mean over all subsamples of the respective size.

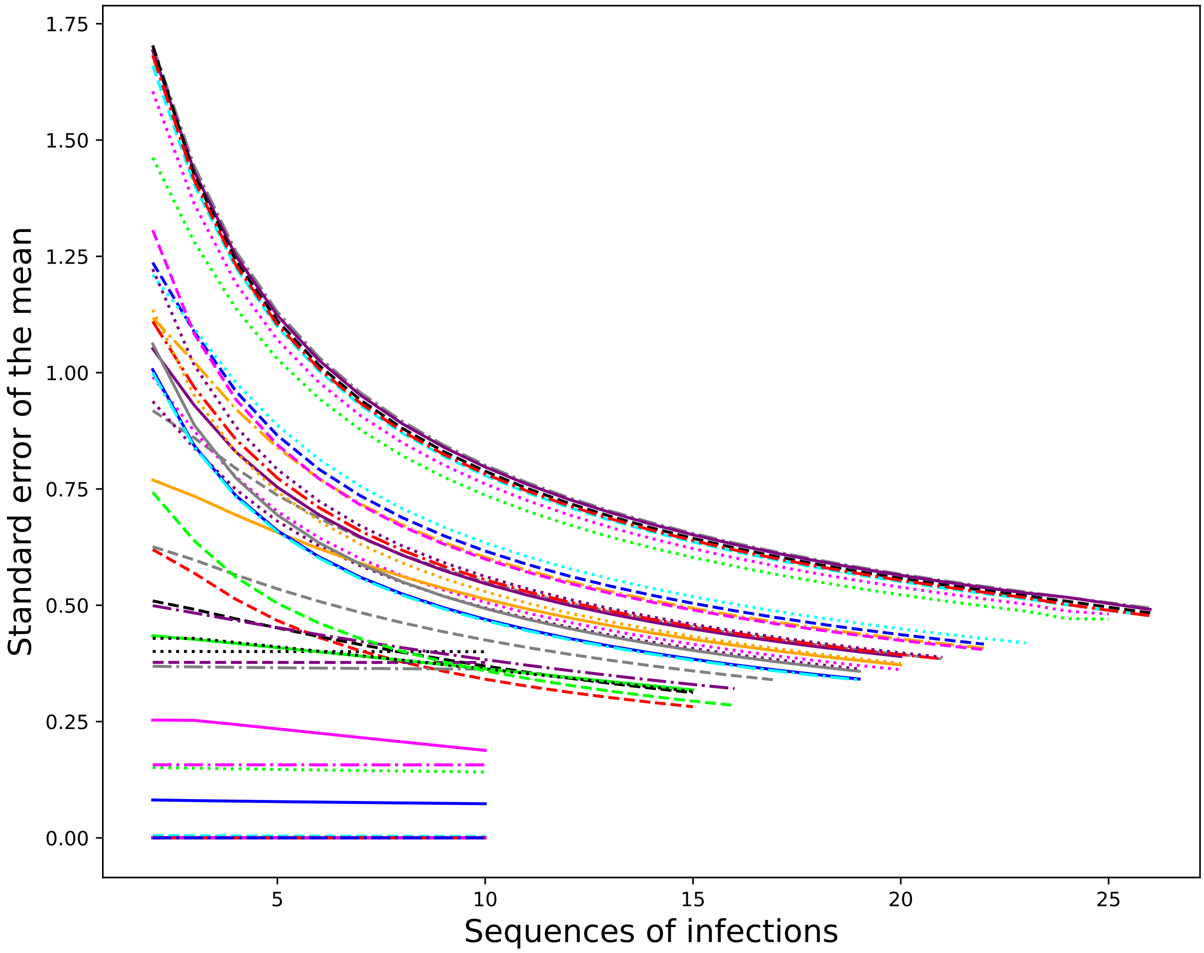


**Figure S5.** The standard error of the mean (as defined in equations S45, S46 and S47) as a function of the sample size. Each line represents a combination of scenario, ciprofloxacin exposure amount and mutation rate. The values are obtained by averaging the standard error of the mean over all subsamples of the respective size.

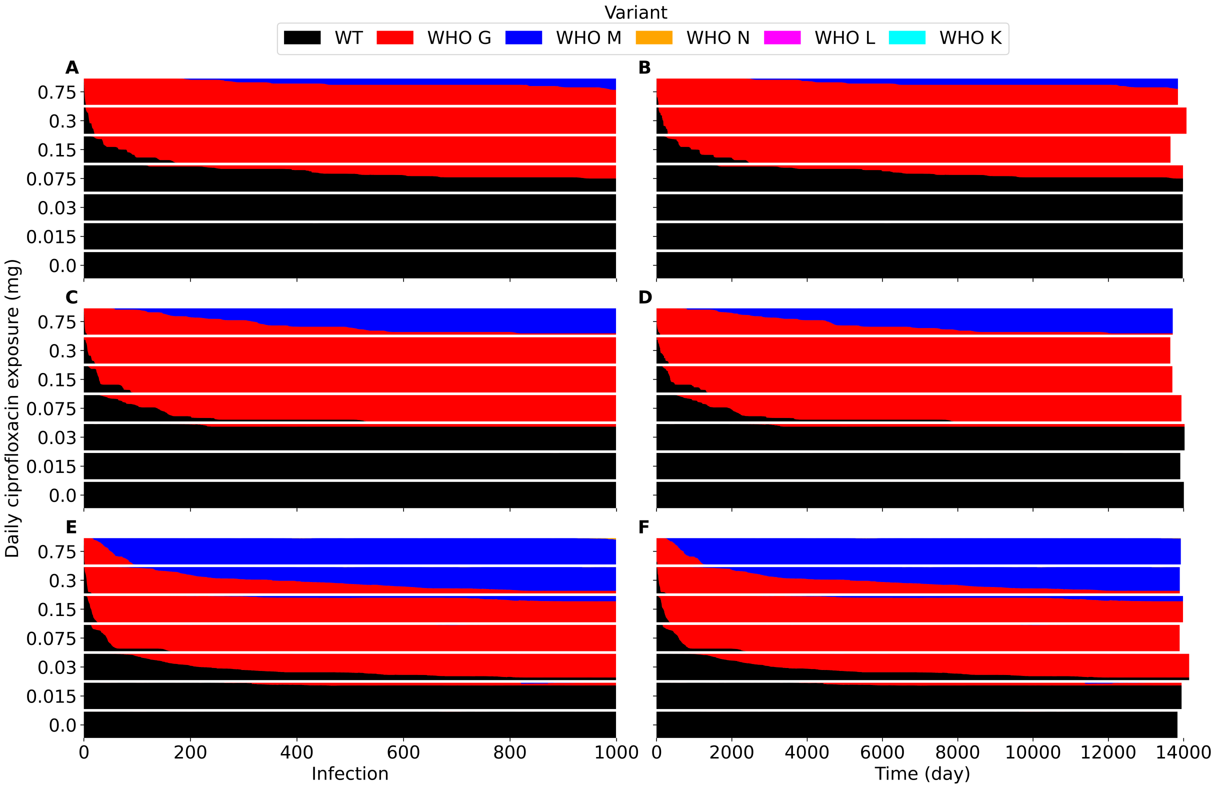


**Figure S6.** Emergence and selection for resistance during a sequence of 1,000 infections under different daily ciprofloxacin exposures. This was simulated without cost of resistance, starting with fully susceptible infections. Each bar shows the distribution of variants across 10 sequences of infections, with the height of each colour indicating the contribution of the corresponding variant. Each row of two panels shows the resistance emergence and selection for a different mutation rate $(10^{-9}$ (**A**, **B**), $10^{-8}$ (**C**, **D**) and $10^{-7}$ (**E**, **F**)). The panels in the left column (**A**, **C**, **E**) show the distribution per infection, while those in the right column (**B**, **D**, **F**) show the distribution per day.

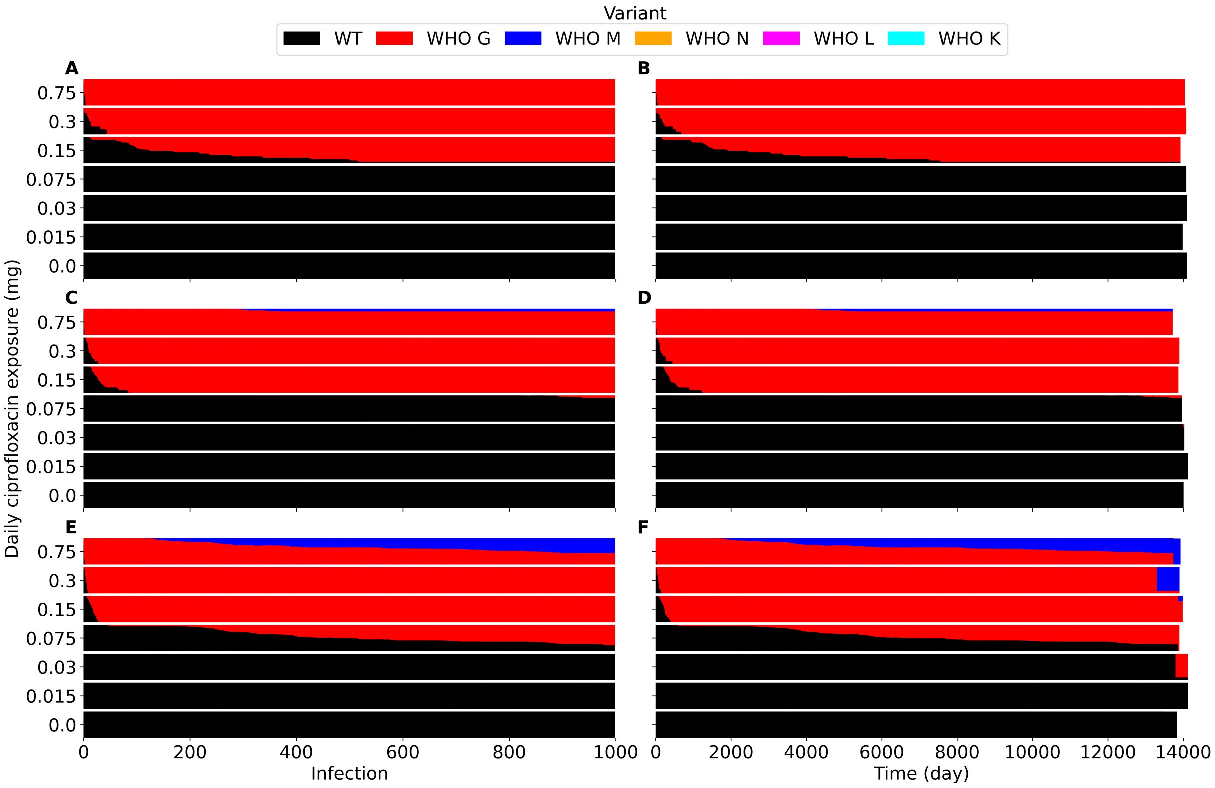


**Figure S7.** Emergence and selection for resistance during a sequence of 1,000 infections under different daily ciprofloxacin exposures. This was simulated with a cost of resistance represented by a 2% growth rate reduction per subsequent resistance mutation, starting with fully susceptible infections. Each bar shows the distribution of variants across 10 sequences of infections, with the height of each colour indicating the contribution of the corresponding variant. Each row of two panels shows the resistance emergence and selection for a different mutation rate $(10^{-9}$ (**A**, **B**), $10^{-8}$ (**C**, **D**) and $10^{-7}$ (**E**, **F**)). The panels in the left column (**A**, **C**, **E**) show the distribution per infection, while those in the right column (**B**, **D**, **F**) show the distribution per day.

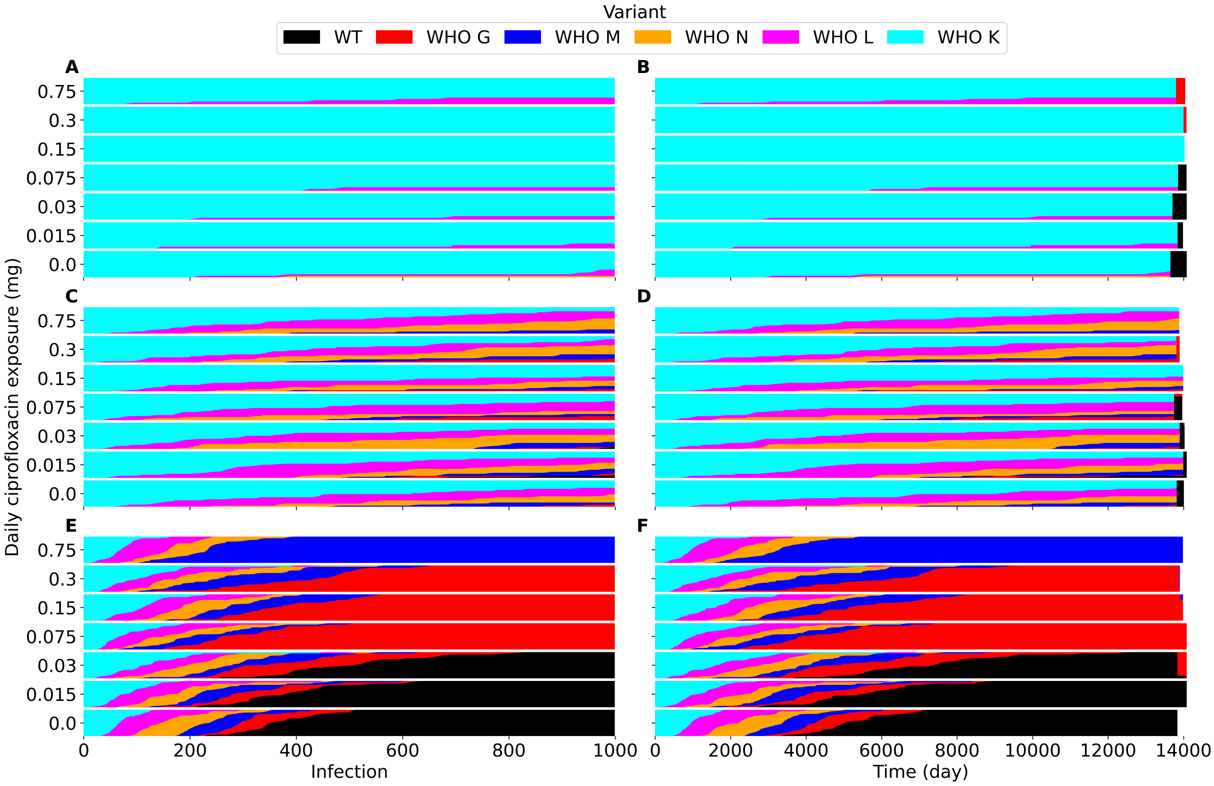


**Figure S8.** Emergence and selection for resistance during a sequence of 1,000 infections under different daily ciprofloxacin exposures. This was simulated with a cost of resistance represented by a 2% growth rate reduction per subsequent resistance mutation and starting with fully resistant infections. Each bar shows the distribution of variants across 10 sequences of infections, with the height of each colour indicating the contribution of the corresponding variant. Each row of two panels shows the resistance emergence and selection for a different mutation rate ($10^{-9}$ (**A**, **B**), $10^{-8}$ (**C**, **D**) and $10^{-7}$ (**E**, **F**)). The panels in the left column (**A**, **C**, **E**) show the distribution per infection, while those in the right column (**B**, **D**, **F**) show the distribution per day.

Sender, R., Weiss, Y., Navon, Y., Milo, I., Azulay, N., Keren, L., Fuchs, S., Ben-Zvi, D., Noor, E., and Milo, R. The total mass, number, and distribution of immune cells in the human body.

*Proceedings of the National Academy of Sciences*, 120(44):e2308511120, 2023.

Shaw, J. and Falkow, S. Model for invasion of human tissue culture cells by neisseria gonorrhoeae. *Infection and immunity*, 56(6):1625–1632, 1988.

Simons, M. P., Nauseef, W. M., and Apicella, M. A. Interactions of neisseria gonorrhoeae with adherent polymorphonuclear leukocytes. *Infection and immunity*, 73(4):1971–1977, 2005.

Simons, M. P., Nauseef, W. M., Griffith, T. S., and Apicella, M. A. Neisseria gonorrhoeae delays the onset of apoptosis in polymorphonuclear leukocytes. *Cellular microbiology*, 8(11):1780–1790, 2006.

Talja, M., Lehtola, A., Salmenperä, M., Ruutu, M., KARONEN, S.-L., and Alfthan, O. Urethral blood circulation during cardiopulmonary bypass. *British journal of urology*, 67(4):381–384, 1991.

Unemo, M., Seifert, H. S., Hook III, E. W., Hawkes, S., Ndowa, F., and Dillon, J.-A. R. Gonorrhoea. *Nature Reviews Disease Primers*, 5(1):79, 2019.

Vance-Bryan, K., Guay, D. R., and Rotschafer, J. C. Clinical pharmacokinetics of ciprofloxacin. *Clinical pharmacokinetics*, 19:434–461, 1990.

Veale, D., Goldner, M., Penn, C., Ward, J., and Smith, H. The intracellular survival and growth of gonococci in human phagocytes. *Microbiology*, 113(2):383–393, 1979.

Wagenlehner, F. M., Kinzig-Schippers, M., Sörgel, F., Weidner, W., and Naber, K. G. Concentrations in plasma, urinary excretion and bactericidal activity of levofloxacin (500 mg) versus ciprofloxacin (500 mg) in healthy volunteers receiving a single oral dose. *International journal of antimicrobial agents*, 28(6):551–559, 2006.

Webb, D., Roberts, D., Williams, J., and Asscher, A. Pharmacokinetics of ciprofloxacin in healthy volunteers and patients with impaired kidney function. *Journal of Antimicrobial Chemotherapy*, 18(Supplement_D):83–87, 1986.
